# Complimentary action of structured and unstructured domains of epsin supports clathrin-mediated endocytosis at high tension

**DOI:** 10.1101/2020.03.27.011437

**Authors:** Jophin G. Joseph, Carlos Osorio, Vivian Yee, Ashutosh Agrawal, Allen P. Liu

## Abstract

Membrane tension plays an inhibitory role in clathrin-mediated endocytosis (CME) by impeding the transition of flat plasma membrane to hemispherical clathrin-coated structures (CCSs). Membrane tension also impedes the transition of hemispherical domes to omega-shaped CCSs, a necessary step before their internalization *via* dynamin-mediated membrane scission. However, CME is not completely halted in cells under high tension conditions. Here we find that epsin, a membrane bending protein which inserts its N-terminus H_0_ helix into lipid bilayer, supports flat-to-dome transition and increases the stability of CCSs at high tension. This discovery is supported by molecular dynamic simulation of the epsin N-terminal homology (ENTH) domain that becomes more structured when embedded in a lipid bilayer. In addition, epsin has an intrinsically disordered protein (IDP) C-terminus domain which induces membrane curvature *via* steric repulsion. Insertion of H_0_ helix into lipid bilayer is not sufficient for stable epsin recruitment as deleting the IDP domain in epsin renders it cytosolic. Epsin’s binding to adaptor protein 2 and clathrin is critical for epsin’s association with CCSs under high tension conditions, supporting the importance of multivalent interactions in CCSs. Together, our results support a model where the ENTH and IDP domains of epsin have complementary roles to ensure CME initiation and CCS maturation are unimpeded under high tension environments.

## Introduction

Clathrin-mediated endocytosis (CME) involves the internalization of cargo by sculpting plasma membrane into 60 −120 nm sized buds, supported by a clathrin protein coat^1^. CME is a well-studied endocytic pathway present in organisms at all developmental stages^1–4^. It plays a critical role in nutrient uptake, intracellular trafficking, and signal transduction^1,3^. CME is a multistep process involving (i) initiation of membrane budding with adaptor proteins and membrane bending proteins^5–7^ (ii) clathrin coat formation^8,9^ (iii) maturation of coated pits^10–12^, and (iv) dynamin-mediated scission of the buds^1,3,7^. Progression of CME involves extensive deformation of flat plasma membrane to Ω-shaped pits^1,12^. Given CME is a mechanical process, membrane tension has been shown to play an inhibitory role during the membrane deformation process preventing the transition from a flat membrane to hemispherical domes^10,11^ and the transition from hemispherical domes to Ω-shaped pits^10–12^. Yet, CME is observed ubiquitously in cells under different membrane tension regimes and this points to the existence of tension-sensitive molecular mechanisms supporting CME^12,13^. Actin-mediated transition of hemispherical domes to Ω-shaped pits at high tension was established by Boulant *et al*^12,14^. However, how membrane-associated proteins aid to overcome the elevated energy barrier needed to initiate budding remains an open question. We hypothesize that endocytic membrane bending proteins possess the ability to sense and counteract membrane tension to facilitate budding at elevated tension.

Epsin/AP180 family is a major family of proteins involved in membrane bending during the initiation of CME^5,15^. Epsin, a prominent member of the family, is believed to insert its N-terminus amphipathic helix (H0 helix in epsin N-terminus homology (ENTH) domain) into the bilayer with a wedging effect to initiate membrane bending^15^. An alternate hypothesis proposed recently posits the C-terminus intrinsically disordered protein (IDP) domain of epsin initiates membrane bending *via* steric crowding^16,17^. *In vitro* studies have shown that insertion of purified ENTH into giant unilamellar vesicles (GUVs) reduces membrane rigidity and area compressibility modulus of the lipid bilayer^18^. Further, recruitment of ENTH softens the bilayer at high tension and initiates tubulation at low tension^19^. An increase in lipid packing defects at high tension may be key in aiding helix insertion at high tension from theoretical studies^20,21^. These evidence points to the existence of a tension-sensitive recruitment mechanism of ENTH domain-containing proteins. However, there is still ambiguity in the exact mechanism of ENTH binding at different tensions, and a lack of experimental evidence for tension-mediated recruitment of epsin to clathrin coat nucleation sites in cells. Here, we used the combination of total internal reflection fluorescence (TIRF) and structural illumination microscopy (SIM) imaging, mechano-manipulation techniques, automated image analysis, and molecular dynamics (MD) simulation to investigate the tension-responsive recruitment of epsin and its stabilization of clathrin-coated structures (CCS) to form clathrin-coated pits (CCPs). We demonstrated that the recruitment of epsin increases with membrane tension in CCSs. Further, using MD simulations and experiments involving epsin mutants, we deciphered the role of H_0_ helix in tension sensing and the molecular mechanism of tension-mediated recruitment of epsin. In addition, we also demonstrate the role of IDP domain of epsin in stabilizing CCSs in high tension environments. Our work establishes a mechanism involving complimentary roles of ENTH and IDP domains of epsin in recruitment into and stabilization of CCSs in a high-tension environment.

## Results

### Overexpression of epsin in cells reduces abortive CCSs at high tension

Previous work from our lab have shown that retinal pigment epithelial (RPE) cells spread on large fibronectin islands (to induce high membrane tension), exhibited an increase in proportion of abortive CCSs and smaller CCPs^22,23^. Using SIM-TIRF super-resolution imaging of RPE cells stably expressing mCherry-clathrin light chain (CLC), we characterized the morphology of CCSs and classified them into three categories: (i) Abortive CCSs, which are coated structures which dissemble before they reach maturation^24–26^, (ii) Stalled CCSs, which are persistent, noninternalizing coated structures in the imaging field^8,12^, and (iii) Productive CCPs, which are coated structures undergo initiation, assembly, and transition to coated-pits followed by membrane scission and internalization. Lifetime analyses of CCSs have shown that abortive CCSs have a lifetime less than 20 s^27–29^ and stalled CCSs have a lifetime above 120 s^8,12,27^. Under SIM-TIRF field, a matured CCS reaches a domed shape and is manifested as a ring (Fig. 1a, shown with red arrow)^30^. The track of a productive CCS shows the evolution of a point signal into a ring structure, subsequently transitioned to a point signal, and finally disappeared from the SIM-TIRF field (Fig. 1a). By comparison, an abortive CCS did not transition from a point signal to a ring structure and disappeared from the SIM-TIRF field within a short time. Stalled CCSs were often clusters of CCSs in the shape of interconnected rings, which remained persistently in the SIM-TIRF field.

**Figure 1.**
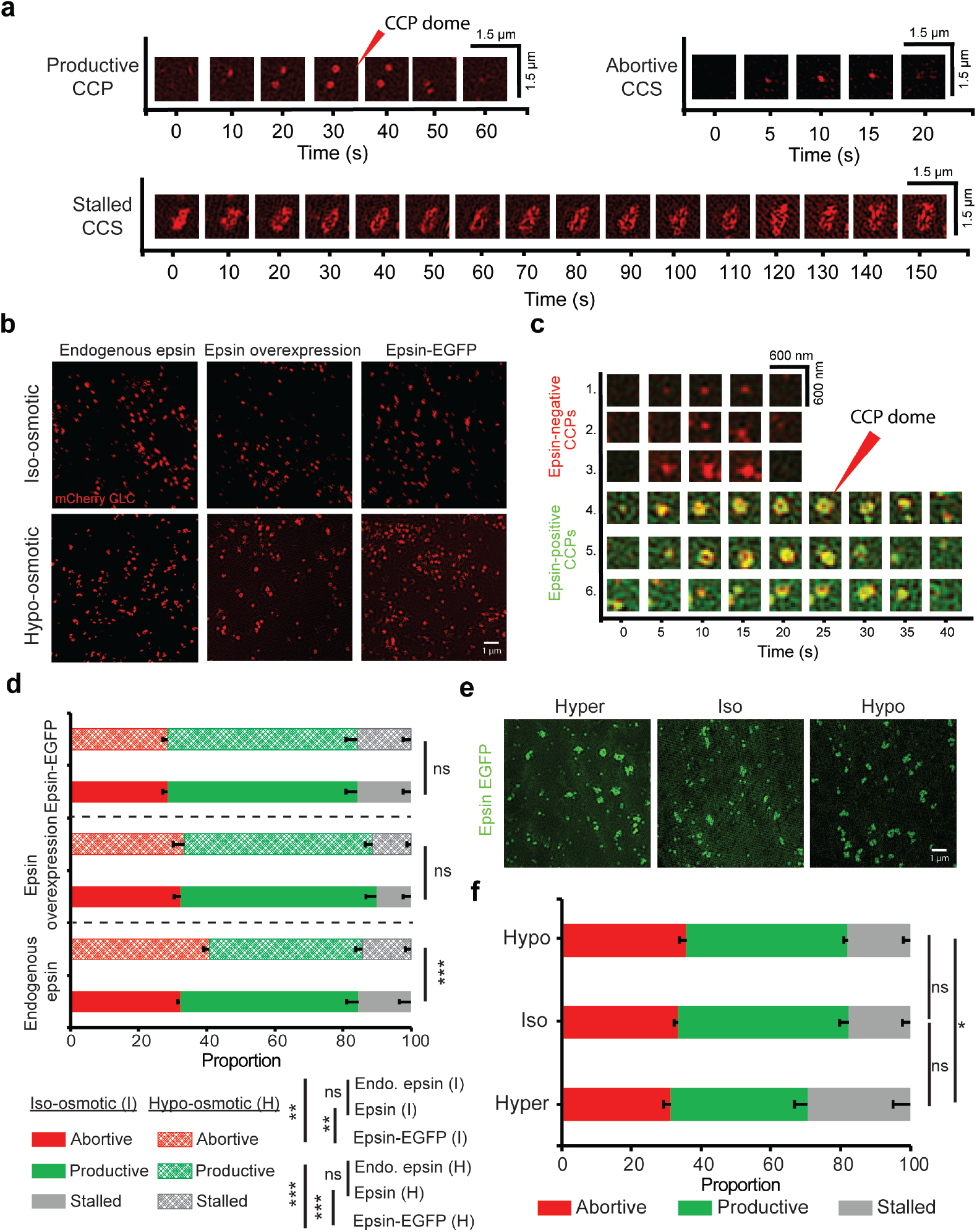
Overexpression of epsin in RPE cells reduces population of abortive CCSs at high tension. **a** Lifetime montage of CCS/CCP (marker: mCherry CLC) classified as abortive, productive and stalled imaged using SIM-TIRF. The formation of a CCS dome (manifested as a ring in SIM-TIRF) is shown with a red arrow. **b** SIM-TIRF images of clathrin in RPE cells expressing epsin at endogenous levels, overexpressing epsin, and overexpressing epsin EGFP under iso-osmotic (control-290 mmol/kg) and hypo-osmotic (220 mmol/kg) conditions. **c** Lifetime montage of CCSs (marker: mCherry CLC) with or without recruitment of epsin (green) using dual color SIM-TIRF. The formation of a CCS dome (manifested as a ring in SIM-TIRF) is shown with a red arrow. **d** Percentage of abortive, productive and stalled CCPs/CCSs in cells expressing endogenous epsin, overexpressing epsin, and overexpressing epsin EGFP under iso-osmotic (control) and hypo-osmotic conditions. **e** SIM-TIRF images of epsin EGFP in RPE cells under hyper- (440 mmol/kg), iso- (290 mmol/kg) and hypo-osmotic (220 mmol/kg) conditions. f. Percentage of abortive, productive, and stalled epsin EGFP structures under hyper-, iso- and hypo-osmotic conditions. For d, the number of CCSs analyzed for endogenous epsin, epsin overexpression, and epsin EGFP overexpression were 26,240 (iso) and 16,918 (hypo), 17,914 (iso) and 14,722 (hypo), 31,776 (iso) and 41,949 (hypo), respectively taken from 6 cells under each condition. For f, the number of epsin EGFP structures analyzed were 31,404 (hyper), 36,536 (iso), and 31,085 (hypo), 31,776 (iso) respectively taken from 6 cells under each condition. The error bars denote standard deviation. NS denotes not significant. *, **, *** represent *p* < 0.01, *p* < 0.001 and *p* < 0.0001, respectively.

RPE cells expressing epsin at endogenous level, overexpressing epsin, and overexpressing epsin EGFP were imaged under iso-osmotic (control - 290 mmol/kg osmolarity) and hypo-osmotic conditions (high tension - 220 mmol/kg osmolarity) using SIM-TIRF (Fig. 1b).

Hyper- and hypo-osmotic shock led to a reduction or increase in cell volume, as evident from 3D cross section of the cells reconstructed from confocal z stacks (Sup Fig 1a). Micropipette aspiration assay revealed that membrane aspiration into micropipette reduced from hyper- to iso- to hypo-osmotic condition (Sup. Fig. 1b-d), similar to what was previously reported^31^. Dual color SIM-TIRF imaging was used to identify co-localization of epsin and clathrin in CCSs (Fig. 1c). The cells were imaged between 5 and 35 minutes after adding the new osmotic media. This is to ensure that the imaging was performed before the cells re-equilibrate their shape^8,32^. Cells under iso-osmotic conditions showed similar proportions of abortive CCSs irrespective of epsin expression level (Fig. 1d). However, overexpressing epsin-EGFP led to a reduction in the percentage of abortive CCSs compared to endogenous expression or overexpression of WT epsin under iso-osmotic condition. On the other hand, when membrane tension was increased under hypo-osmotic conditions, cells expressing endogenous level of epsin showed an appreciable increase in abortive CCSs, consistent with previous findings^22^. In contrast, overexpression of WT epsin or epsin EGFP both maintained the same percentages of abortive CCSs under hypo-osmotic condition compared to iso-osmotic condition (Fig. 1d). Interestingly, cells overexpressing epsin EGFP had a higher percentage of stalled CCSs compared to cells overexpressing WT epsin. As a result, overexpression of WT epsin had a larger fraction of productive CCPs compared to epsin EGFP overexpression. Further, membrane-associated structures with epsin EGFP were imaged at hyper- (low tension - 440 mmol/kg osmolarity), iso- (control - 290 mmol/kg osmolarity) and hypo- (high tension - 220 mmol/kg osmolarity) osmotic conditions (Fig. 1e). As membrane tension increased, the percentage of abortive structures with epsin recruitment increased only slightly (Fig. 1f). However, at low tension, epsin recruitment resulted in CCS clustering and stalling (Fig. 1f). This stalling and clustering is consistent with *in vitro* observation of epsin tubulation at low tension in GUVs^16–19^. Even though clusters of CCSs which are stalled remained in the TIRF field during the acquisition period, they showed dynamic rearrangement of CCS rings including merging and splitting of rings (Sup. movie 1). Some clusters showed disappearance of CCSs from the boundary of the stalled clusters (Sup. movie 1).

### Epsin recruitment increases as resting membrane tension increases

Epsin is an endocytic adaptor protein recruited to the plasma membrane during initiation of CME to aid in membrane curvature generation^15^. The ability for pits to maintain stable curvatures until they reach maturation is essential for successful completion of a CCS lifecycle^12^. Membrane tension is shown to counteract the stabilization of coated pits by increasing the energy requirement to form a pit^11^. To investigate whether epsin recruitment can counteract varying membrane tension, we manipulated resting or acute membrane tension of RPE cells co-expressing epsin EGFP and mCherry CLC by controlling cell spreading or by applying osmotic shock, respectively.

Resting tension of an adherent cell is related to the membrane tension exerted when the cell is fully spread^33,34^. The resting membrane tension of the cells was controlled by restricting cell spreading on microcontact-printed fibronectin islands of area 625 μm^2^ or 1024 μm^2^ (Fig. 2a). Cells spread on a larger area experienced higher membrane tension^22,33^. Using dual color TIRF imaging and an automated algorithm^27^, we detected and tracked CCSs and quantified the lifetime, composition (whether both epsin and CLC are present or not), and intensity of fluorescently tagged epsin (EGFP) and CLC (mCherry) in CCSs. The intensity traces of CCSs belonging to 60-78 s lifetime cohort showed that the intensity of EGFP epsin is higher in highly spread cells, suggesting epsin recruitment increases with an increase in resting tension (Fig. 2b). Interestingly, epsin intensity is already elevated at time 0 when clathrin detection began, suggesting epsin EGFP recruitment may precede clathrin recruitment. Further, the average plateau intensity (i.e., the maximum intensity during a coated pit’s lifetime) of epsin in other lifetime cohorts (from 10-18 s to 80-98 s) also showed a higher recruitment of epsin to CCSs in more spread cells in comparison with the less spread cells (Fig. 2c). In contrast, the intensity of mCherry CLC in the 80-98 s lifetime cohort and other lifetime cohorts had a small reduction as resting tension increased (Sup. Fig. 2a and 2b), in agreement with a previous finding^22^. Together, these results indicate that epsin recruitment per clathrin in CCSs increases with increasing resting tension.

**Figure 2.**
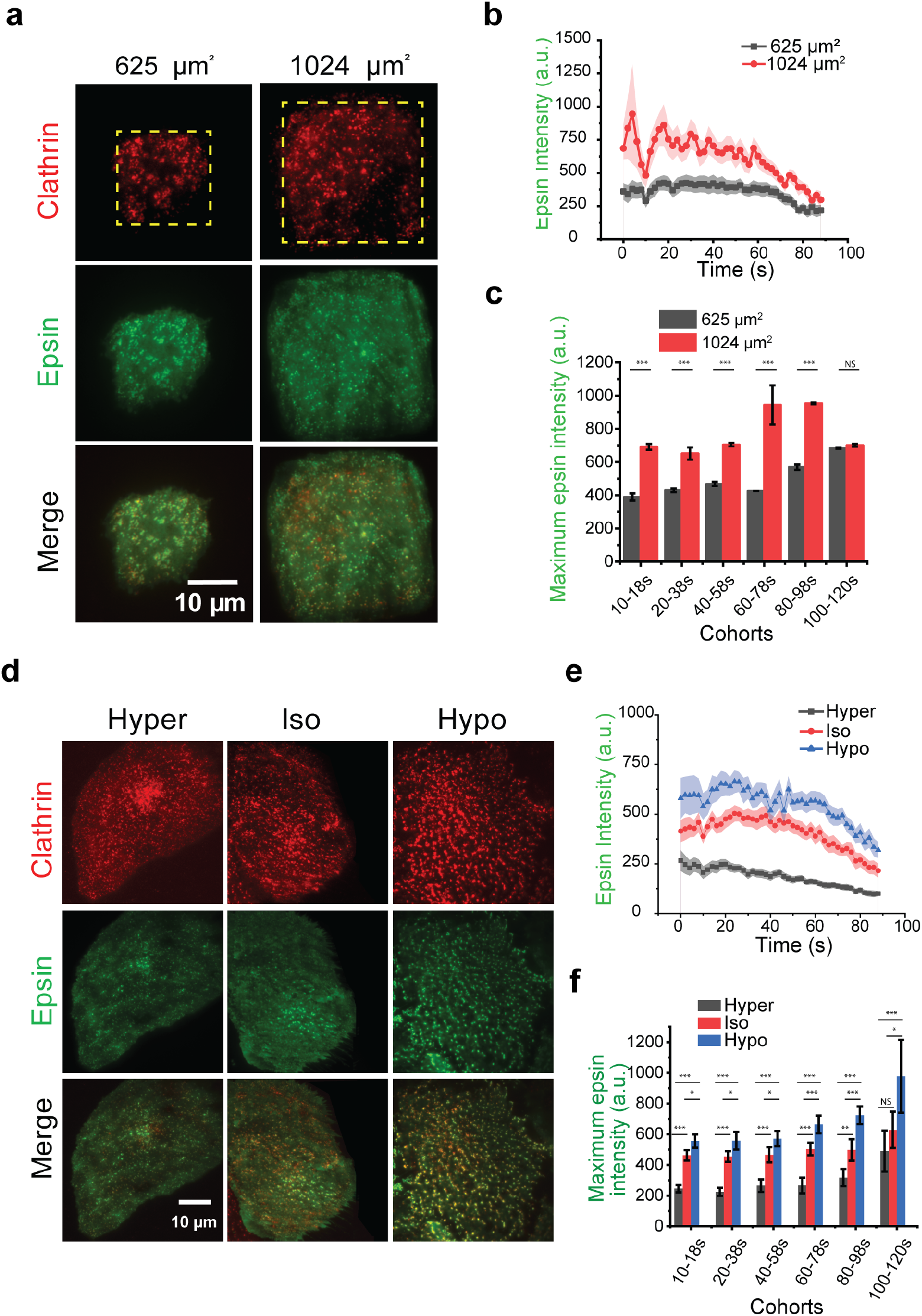
Increased tension increases recruitment of epsin into CCSs. **a** RPE cells expressing epsin EGFP and mCherry CLC spread on fibronectin square islands of size 25 and 32 μm. The resting membrane tension is higher for 32 μm islands. **b** The average intensity trace of epsin on 25 μm (black) and 32 μm (red) islands for CCSs with 60 -78 s lifetime. **c** The average plateau intensity of epsin across different CCS lifetime cohorts for cells on 25 μm (black) and 32 μm (red) islands. **d** Fluorescence images of clathrin and epsin of RPE cells expressing epsin EGFP under different osmotic conditions. **e** The average plateau intensity of epsin in different osmotic media for CCSs with 60 -78 s lifetime. **f** The average plateau intensity of epsin across different CCS lifetime cohorts. For b, c, N_cells_ for 25 μm square and N_cells_ for 32 μm square are 18 and 19, respectively. For e, f, the number of cells for hyper-, iso-, and hypo-osmotic conditions were 19, 19, and 21, respectively. The error bars denote standard error. NS denotes not significant. *, **, *** represent *p* − 0.01, *p* < 0.001 and *p* < 0.0001, respectively.

### Epsin recruitment increases as acute membrane tension increases

To investigate whether epsin shows a similar recruitment characteristic in response to acute tension changes, we used osmotic shock to induce acute tension changes in RPE cells. When imaged by TIRF microscopy, epsin puncta appear to be brighter under hypo-osmotic conditions (Fig. 2d). The intensity traces of CCSs belonging to 60-78 s lifetime cohort showed an increase in intensity of epsin EGFP when the osmolarity of the media was reduced (Fig. 2e). The average plateau intensity of epsin in other lifetime cohorts (from 10-18 s to 100-120 s) was also higher with a decrease in media osmolarity (Fig. 2f), a result similar to what we have observed from the resting cell tension experiments. Our osmotic shock experiments suggest an increase in recruitment of epsin accompanies an acute increase in tension. The intensity of mCherry CLC in CCSs in epsin-expressing cells, belonging to 60-78 s lifetime cohort, showed a small increase with the acute increase in tension (Sup. Fig. 2c). CLC plateau intensity in epsin EGFP-recruited CCSs were the same between hyper- and iso-osmotic conditions but significantly increased for hypo-osmotic condition (Sup. Fig. 2d). This result is different from the resting cell tension experiments where we observed reduced clathrin intensity, suggesting an acute tension increase has a different effect on clathrin assembly. Nevertheless, clathrin recruitment into CCSs in cells without epsin overexpression (Sup. Fig. 3a) was lower when cells were subjected to an acute increase in tension (Sup. Fig. 3b). Thus, the recruitment of epsin could promote clathrin recruitment when membrane tension increases acutely.

### ENTH domain containing H_0_ helix supports tension-mediated nucleation of epsin at CCS sites

The tension response by epsin to changes in resting tension or acute tension variation is unique from other membrane-associated proteins. Typically, elevated tension reduces the recruitment of proteins to the membrane^22,35^. Paradoxically, epsin shows an increase of recruitment in response to elevated tension. Since the H_0_ helix in the N-terminus of epsin can insert into the bilayer upon membrane binding^15^, we hypothesize that this helical insertion contributes to the unique tension response of epsin. To test this hypothesis, we first removed the ENTH domain of epsin, which is the structured N-terminus region of epsin that contains multiple alpha helices, including the H_0_ helix. The removal of ENTH domain of epsin did not render it cytosolic (Fig. 3a), suggesting other parts of epsin can still target it to CCSs. However, it appeared to decrease the number of puncta compared to full-length epsin as membrane tension was increased (Fig. 3a). Further, as tension increased, overexpressing full-length epsin led to an increase in CCS initiation density (Fig. 3b). This result implies that epsin can sense an increase in membrane tension and respond to it dynamically by increasing CCS nucleation. In contrast, the initiation density of epsin ΔENTH puncta remained unchanged with an increase in membrane tension (Fig. 3b). This points to ENTH domain playing a role in tension sensitivity of epsin and mediating epsin membrane binding at high membrane tension.

**Figure 3.**
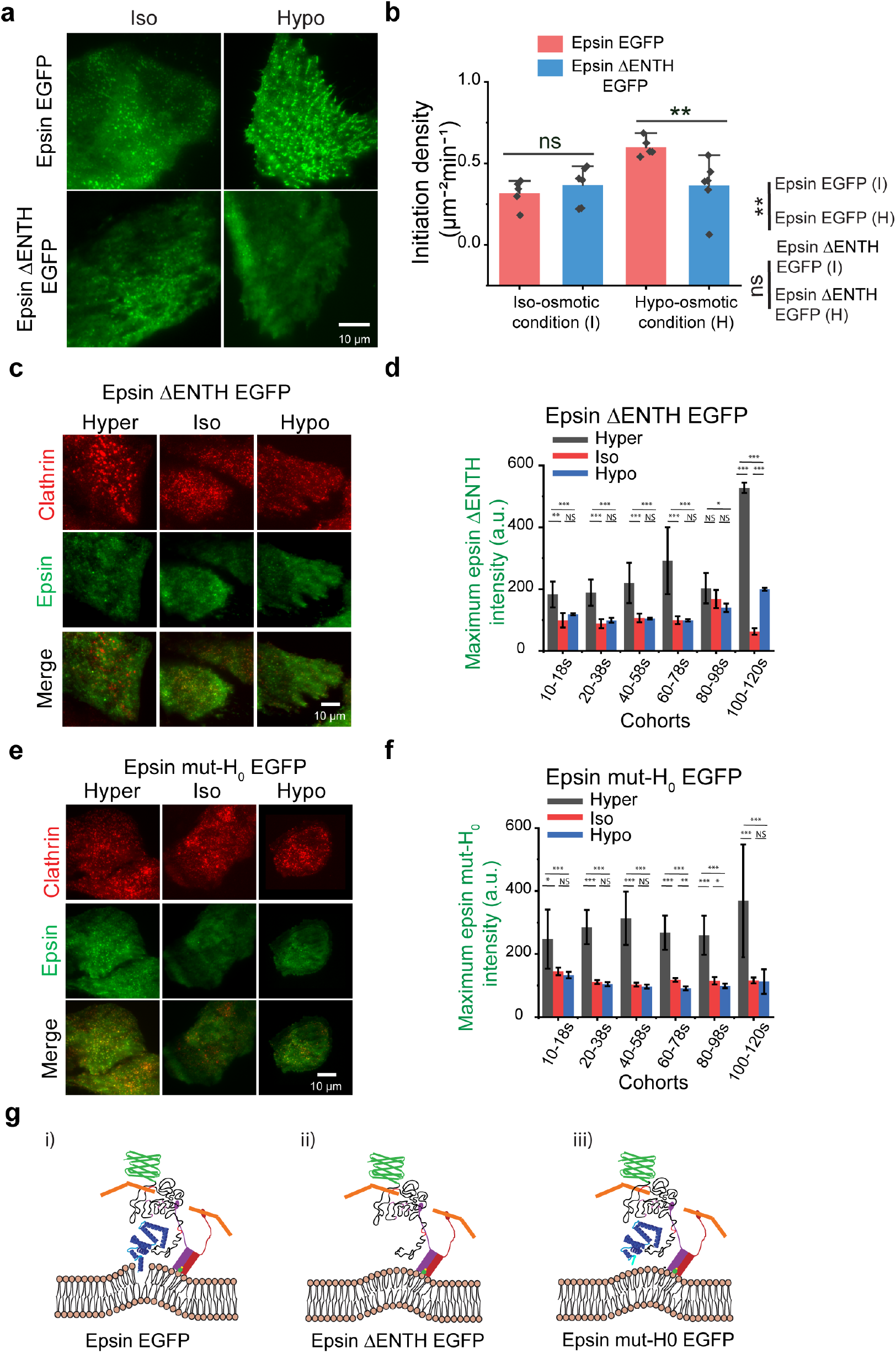
ENTH domain containing H_0_ helix supports tension-mediated nucleation of epsin at CCS sites. **a** Fluorescence images of epsin, epsin ΔENTH at control (iso-) and high-tension(hypo-osmotic conditions) **b** Initiation density of epsin puncta in epsin EGFP (salmon) expressing and epsin ΔENTH (blue) expressing cells under control (iso-osmotic) and high-tension (hypo-osmotic) conditions. **c** Representative fluorescence images of clathrin and epsin of RPE cells expressing epsin ΔENTH EGFP under different osmotic conditions. **d** The average plateau intensity of epsin ΔENTH in different CCS lifetime cohorts in different osmotic conditions. **e** Representative fluorescence images of clathrin and epsin of RPE cells expressing epsin mut-Ho EGFP under different osmotic shock conditions. **f** The average plateau intensity of epsin mut-Ho in different CCS lifetime cohorts in different osmotic conditions. **g** Proposed model for (i) epsin EGFP, (ii) epsin ΔENTH EGFP, and (iii) epsin mut-Ho EGFP nucleation at CCS sites. For b, N_cells_ expressing epsin EGFP under iso- and hypo-osmotic conditions were 9 each, and N_cells_ expressing epsin ΔENTH under iso-and hypo-osmotic conditions were 9 each. The N_cells_ expressing epsin ΔENTH EGFP for hyper-, iso-, and hypo-osmotic conditions in d. were 20, 20, and 20, respectively. The N_cells_ expressing epsin mut-Ho for hyper-, iso-, and hypo-osmotic conditions in f. were 19, 24, and 18, respectively. The error bars denote standard error. NS denotes not significant. *, **, *** represent *p* < 0.01, *p* < 0.001 and *p* < 0.0001, respectively.

To further delineate the effect of ENTH domain on tension-mediated recruitment of epsin, we analyzed the intensity of epsin ΔENTH in CCSs in cells co-expressing epsin ΔENTH EGFP and mCherry CLC (Fig. 3c). The average plateau intensity of epsin ΔENTH EGFP in CCPs across multiple lifetime cohorts reduced in response to osmotic shock-induced membrane tension increase (Fig. 3d). Interestingly, decreasing membrane tension using hyper-osmotic shock drastically increased epsin ΔENTH EGFP in CCSs, but how this occurred was not clear. Removal of ENTH domain abrogated the elevated epsin recruitment to CCSs at high tension. However, the average plateau intensity of clathrin in CCSs with epsin ΔENTH EGFP reduced or remained comparable between iso-osmotic and hypo-osmotic conditions (Sup. Fig. 4a). This hinted the possibility that ENTH domain may be dispensable for epsin-mediated stabilization of clathrin in CCSs.

To understand how removal ENTH render epsin unable to sense tension, We next replaced the residues in ENTH which form H_0_ helix upon binding PIP_2_ ^36^ to alanine. These mutations prevent the formation of H_0_ helix when epsin binds to lipid bilayer and blocks amphipathic helix insertion. Dual-color TIRF imaging (Fig. 3e) and downstream analysis of CCPs with mut-H0 epsin co-localization showed a reduction in mut-H0 epsin recruitment across different lifetime cohorts from an acute tension increase, whereas a significant increase in mut-H0 epsin recruitment was again observed when tension was reduced (Fig. 3f). This confirmed that H_0_ helix imparts the tension sensitivity to epsin. This is a significant result, as we showed that ENTH domain with amphipathic insertion can sense membrane tension in addition to the already known ability to detect membrane curvature^36,37^ and dynamically alter the recruitment pattern of epsin to CCSs. Similar to epsin ΔENTH EGFP, the average plateau intensity of clathrin in CCPs with epsin mut-H0 EGFP reduced slightly or remained in the comparable range as membrane tension increased acutely (Sup. Fig. 4b). This suggests that epsin-mediated stabilization of clathrin in CCPs can occur without amphipathic insertion of H_0_ into the bilayer. Finally, we believe that endocytic binding domains in C-terminus region of epsin helps its binding to CCSs in the absence of ENTH domain or amphipathic insertion. Based on this assumption, we propose a model for epsin binding for full-length (Fig. 3g (i)), ΔENTH (Fig. 3g (ii)), mut-H0 (Fig. 3g (iii)).

### Atomistic insights into ENTH-membrane interactions

We performed MD simulations to quantify the interactions of ENTH domain with a lipid membrane made of 1-palmitoyl-oleoyl-sn-glycero-phosphocholine (POPC) lipids and a single PIP2 lipid using CHARM-GUI. The ENTH domain was placed onto the membrane. This led to an instantaneous interaction of the H_0_ helix with PIP_2_ and the subsequent insertion of the H_0_ helix into the membrane. The ENTH domain was then pulled away from the protein. As a consequence, the inserted H_0_ helix first transitioned to an adsorbed state and finally left the membrane and went into the solution. The three stages are shown in Fig. 4a. It is notable that the secondary structure of the H_0_ helix undergoes transformation based on the degree of lipid interactions. Fig. 4b shows the secondary structure analysis as a function of the three stages of the H_0_ helix. In the inserted state, the H_0_ has an alpha-helix structure (pink color). In the adsorbed state with reduced lipid interactions, the H_0_ begins to become disordered from the N-terminus (teal color). In the final state when H_0_ is in the solution, the alpha-helix domain shrinks further transforming into the disordered domain (teal color). The extent of helicity present in the H_0_ helix in the three states is summarized in Fig. 4c.

**Figure 4:**
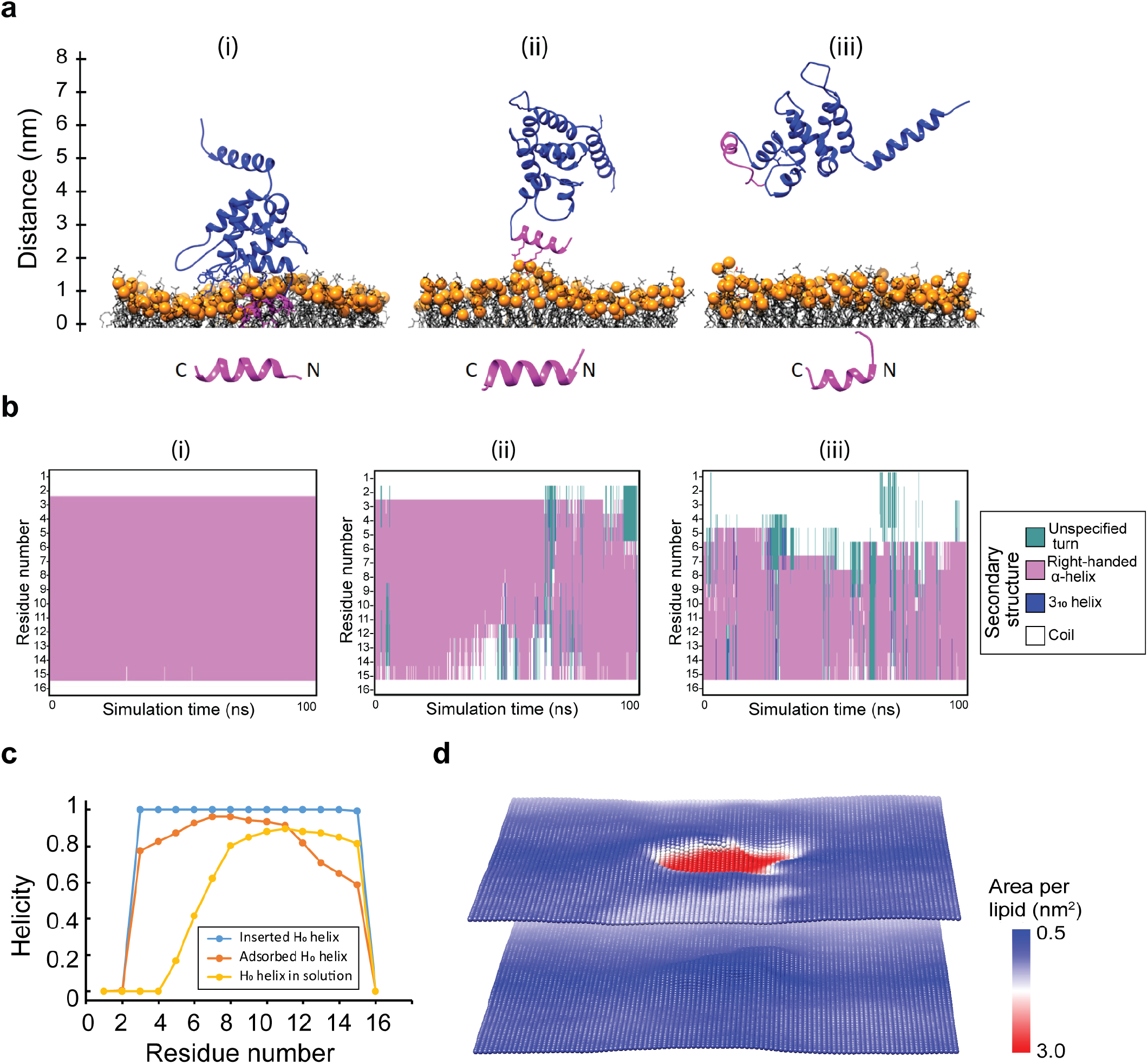
Atomistic insights into ENTH-membrane interactions. **a** The ENTH domain interacting with POPC membrane with a single PIP_2_ lipid. The H_0_ helix shown in magenta is in the inserted, adsorbed and solvent states. **b** The secondary structure analysis of each residue of H_0_ in the three states shown in (a). **c** The summary of the analysis in (b). The helicity of H_0_ decreases as it undergoes reduced interactions with the membrane. The protein-lipid interactions stabilize the alpha-helix structure of Ho. **d** The area per lipid plot corresponding to the inserted state shown in Fig. (a). The POPC lipids have a typical area of 0.64 nm^2^. Presence of H_0_ displaces lipids in the top leaflet. As a result, the effective area per lipid increases in the Ho-occupied domain (red region).

Fig. 4d shows the areal footprint of the H_0_ helix inside the membrane. The plot shows area per lipid in the two leaflets of the membrane. The POPC lipid area is around 0.64 nm^2^ (blue color). Because of the H_0_ helix insertion, the lipids are moved out of the Ho-occupied domain. This displacement of lipids effectively increases lipid-lipid separation, which in turn results in an increase in the area per lipid (red color). Since the protein sits primarily in the top leaflet, the change in area per lipid is minimal in the bottom leaflet. This areal footprint plot suggests a potential mechanism for tension sensitivity exhibited by epsin. A single H_0_ helix occupies an area of 2 nm^2^ (red region) and displaces lipids in the membrane. If the membrane has zero resting tension and the lipids are allowed to move freely, there would be no energetic advantage to displacing the lipids. However, if the membrane has a non-zero resting tension (*σ*), the displacement of lipids would be associated with an energetic incentive of –*σΔA*, where ΔA is the area occupied by the H_0_ helix. This idea is similar to the notion that explains the tension sensitivity of mechanosensitive channels in bacterial membranes^38,39^.

### Interfering the formation of H_0_ helix inhibits early recruitment of epsin to CCPs

The earlier observation that the recruitment of epsin EGFP prior to the arrival of clathrin and the importance of H_0_ helix insertion prompted us to ask whether masking the ENTH domain might alter epsin recruitment to CCPs. Here we considered epsin with EGFP tagged to the C-terminus and EGFP tagged on the N-terminus (masking the ENTH domain) (Sup. Fig. 5a), in addition to epsin ΔENTH EGFP and epsin mut-Ho EGFP mutants, and examined kymographs of different epsin mutants. Using CCP tracks with detection that started at first significant clathrin signal (Fig. 5a), we determined the order of arrival of epsin with respect to clathrin. Epsin with

**Figure 5.**
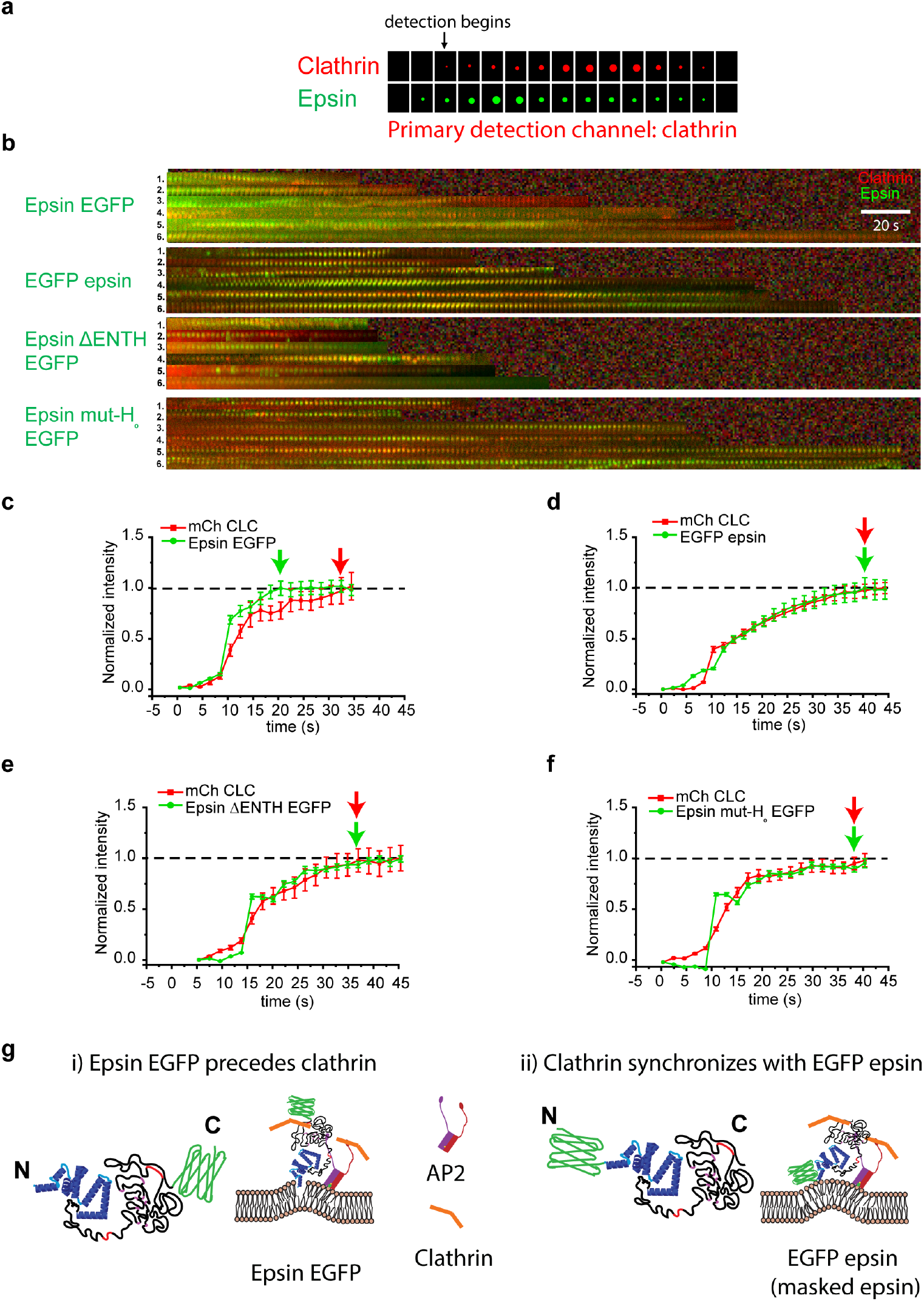
Epsin is recruited prior to clathrin to CCS nucleation sites by amphipathic helix insertion. **a** Schematic of dual-channel detection depicting a scenario in which epsin appears prior to clathrin. **b** Kymographs of CCSs with epsin EGFP, EGFP epsin, epsin ΔENTH EGFP and epsin mut-H0 EGFP (primary detection channel: clathrin). Normalized recruitment of epsin (green) and clathrin (red) (primary detection channel: epsin) for the 60-78s lifetime cohorts until they reach maximum intensity for (**c**) epsin EGFP, (**d**) EGFP epsin, (**e**) epsin ΔENTH EGFP, and (**f**) epsin mut-H0 EGFP. **g** Proposed model for epsin recruitment with (i) EGFP in C-terminus and (ii) EGFP in N-terminus. N_cells_ for d, e, f, and g were 19, 18, 20, and 24, respectively. The error bars denote standard error.

EGFP attached to the C-terminus arrived at CCPs prior to clathrin, as the kymograph showed more green fluorescence at the beginning of the lifetime track (Fig. 5b panel 1), pointing to the early recruitment of epsin prior to clathrin and consistent with what we observed in Fig. 2b and 2f. In contrast, the masked ENTH showed synchronous recruitment with clathrin (Fig. 5b panel 2). Similarly, both epsin ΔENTH EGFP and epsin mut-H0 EGFP helix mutants showed synchronous recruitment with clathrin (Fig. 5b panel 3 and 4). These data were further quantified by plotting the normalized intensities of clathrin and epsin until they reached their maximum intensity after performing CCP detection and tracking with epsin as primary channel. Normalized epsin EGFP recruitment curve was above the mCherry clathrin recruitment curve, pointing to early recruitment of epsin (Fig. 5c). In addition, epsin EGFP reached maximum recruitment prior to mCherry CLC (shown with green and red arrow). However, cells expressing epsin with masked or deleted ENTH domain or with mutated H_0_ helix showed epsin recruitment in synchrony with mCherry CLC (Fig. 5 d-f). Based on these results, we suggest that epsin EGFP recruitment to membrane precedes clathrin (Fig. 5g case i), whereas masking ENTH domain with EGFP disrupts or delays the formation of H_0_ amphipathic helix insertion into the bilayer (Fig. 5g case ii). This is also supported by the synchronous recruitment of epsin mut-H0 EGFP and epsin ΔENTH with clathrin, as they cannot form H_0_ helix upon binding to the membrane.

### Bi-directional stabilization of epsin recruitment to CCSs and curvature of CCS domes is mediated by the IDP domain

Removal of ENTH domain did not render epsin cytosolic. Hence, to determine which domains are responsible for the stabilization effect of epsin, we progressively removed binding domains from epsin within the IDP domain (Sup. Fig. 5b). Removal of clathrin binding domain (CBD) 1 (LMDLADV) or CBD2 (LVDLD) did not affect epsin recruitment into CCSs (Sup. Fig. 6 (i) and (ii))^40^. Simultaneous removal of CBD1 and CBD2 also did not inhibit formation of epsin puncta (Sup. Fig. 6 (iii)), nor was the number of epsin puncta in these mutants affected by an acute increase in membrane tension *via* osmotic shock (Sup. Fig. 6 (i) to (iii), top and bottom panel). However, removal of DPW repeat motifs, which bind to AP2 subunit along with CBD1 and CBD2, render epsin cytosolic (Sup. Fig. 6 (iv)). Using a transferrin uptake assay, we showed that overexpression of epsin mutants did not affect cargo recruitment and internalization *via* CME (Sup. Fig. 7), suggesting that mutant epsins did not impact CME. Consistent to earlier finding, removing the unstructured IDP domain of epsin containing all the endocytic binding sites also resulted in epsin being cytosolic, and this is true under all tension conditions (Fig. 6a). This implies that ENTH, which is the structured domain of epsin containing H_0_ helix, alone cannot stabilize epsin recruitment to CCSs, but it requires the AP2 and clathrin binding sites in the C-terminus unstructured region of the protein. This is a significant result as we have shown that ENTH domain mediates tension-dependent nucleation of epsin and *in vitro* studies have shown ENTH domain alone is recruited to pre-curved lipid bilayers containing PIP_2_^15,37^. To further confirm the requirement of AP2 and clathrin binding to stabilize epsin recruitment to plasma membrane, we used shRNA knockdown of clathrin heavy chain (CHC) or α-adaptin in AP2 subunit. Knocking down CHC or α-adaptin that blocks the formation of CCSs also inhibited the formation of epsin puncta (Sup. Fig. 8). This behavior was observed in full-length epsin, epsin ΔENTH and epsin ΔIDP expressing cells. This provides support that binding of epsin to CHC and AP2 is necessary for stable recruitment of epsin and the ENTH domain alone cannot support stable plasma membrane recruitment (Fig. 6b).

**Figure 6.**
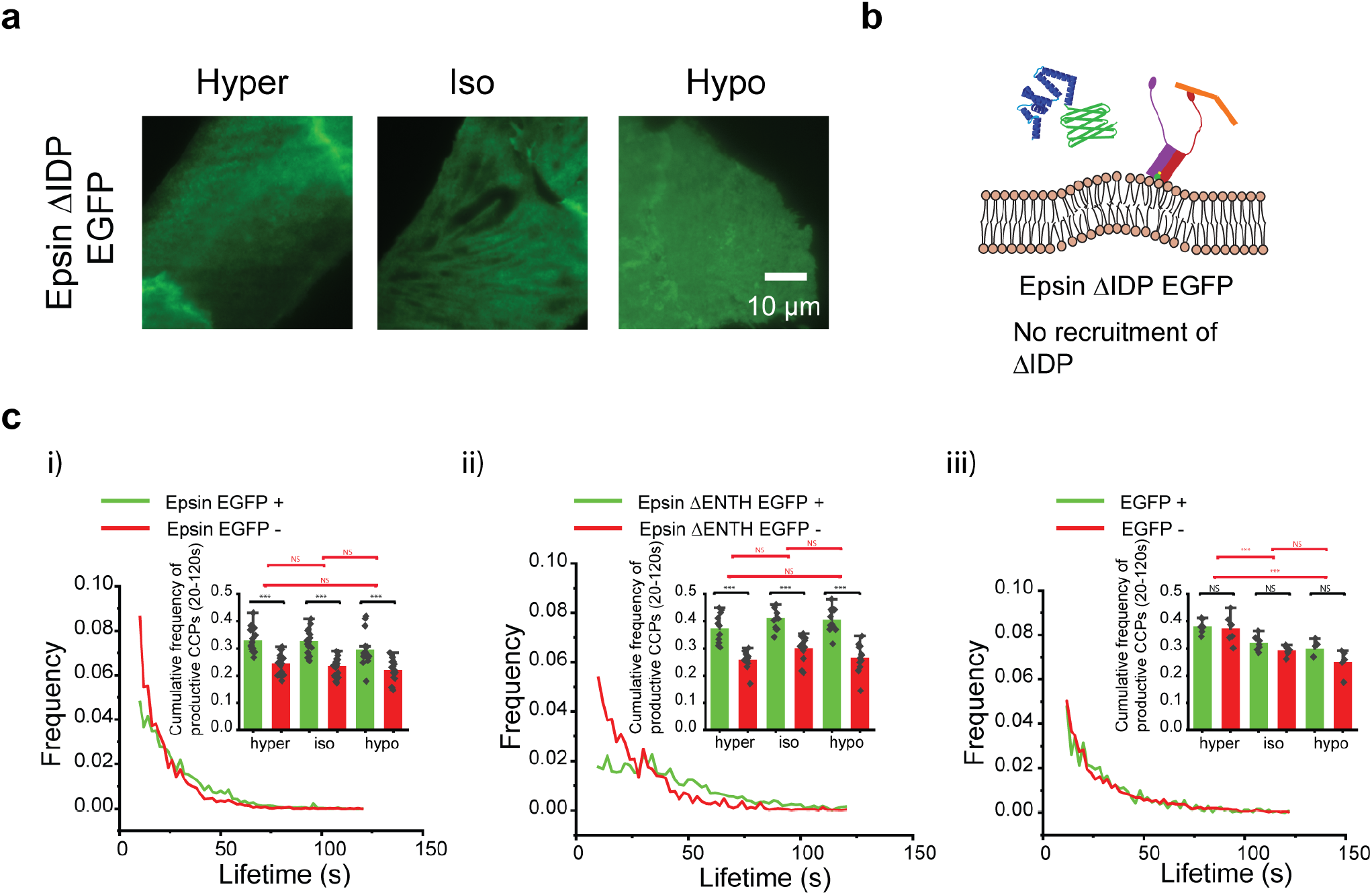
Bi-directional stabilization of epsin recruitment in CCSs and curvature of CCS domes to form productive CCSs is mediated by IDP domain. **a** Fluorescence images of epsin ΔIDP at different tension conditions (hyper-, iso- and hypo-osmotic conditions). **b** Proposed model of interaction between CCS and epsin ΔIDP. **c** The lifetime distribution of CCSs with (green) or without (red) epsin EGFP for iso-osmotic condition for (i) full length epsin with C-terminus EGFP, (ii) epsin ΔENTH EGFP, and (iii) EGFP (control). Inset shows cumulative frequencies of mature CCSs with lifetimes 20-120 s for the different osmotic conditions indicated. The N_cells_ expressing epsin EGFP for hyper-, iso-, and hypo-osmotic conditions in c. were 19, 19, and 21, respectively. The N_cells_ expressing ΔENTH EGFP epsin for hyper-, iso-, and hypo-osmotic conditions in c. were 20, 20, and 20, respectively. The N_cells_ expressing EGFP for hyper-, iso-, and hypo-osmotic conditions in c. were 20, 19, and 16, respectively. The error bars denote standard error. NS denotes not significant. *, **, *** represent *p* < 0.01, *p* < 0.001 and *p* < 0.0001, respectively.

Productive CCPs (lifetime 20-120 s) internalize into cytoplasm without prematurely dissembling or stalling on the membrane. Our SIM-TIRF data showed that overexpression of epsin supports the maturation of CCSs to productive CCPs. Furthermore, dual color SIM-TIRF data showed that the presence of epsin tagged with EGFP preferentially supports the formation of productive CCPs (shown by the preferential formation of as rings when epsin is present (green) in Fig. 1c), suggesting that epsin may stabilize productive CCPs. To determine the effect of the epsin recruitment on the stability of CCSs, we classified CCS tracks as with or without co-localization with epsin. We found that the fraction of productive CCPs in a cell is higher in epsin-containing CCPs (Fig. 6c (i)). We also found that there was no statistically significant change in the fraction of productive CCPs between hyper-, iso- and hypo-osmotic-treated cells (Fig. 6c (i) inset). Similarly, cells having CCSs with epsin ΔENTH recruitment also showed an increase fraction of productive CCPs (Fig. 6d (ii)), again independent of membrane tension. As a control, we looked at whether cells expressing EGFP had the same fraction of productive CCPs in different tension conditions. For this analysis, we used the false positive puncta detection from the secondary channel (EGFP) to classify pits as EGFP-positive (EGFP +) and EGFP-negative (EGFP -). Comparison of the fractions of productive CCPs which are EGFP + and EGFP-showed no difference between them (Fig. 6d (iii)), unlike epsin-positive CCSs in epsin or epsin ΔENTH expressing cells which showed significant increases in the population of productive CCPs. As tension increased, the fraction of productive CCPs decreased for cells expressing only EGFP, consistent with the previous finding from our lab^22^. Altogether, these findings show that epsin support the formation of productive CCPs. With the help of endocytic binding sites in the IDP domain that confers steric crowding, epsin is able to stabilize curvature of CCS domes. ENTH domain, however, does not play an active role in providing this stabilization effect. This result is significant as it shows a bi-directional stabilization of epsin recruitment by binding to AP2 and CHC and stabilization of productive CCPs due to binding of epsin to AP2 and CHC, both mediated by IDP domain of epsin.

## Discussion

Prior experimental evidence points to actin dynamics in supporting the transition of CCSs from hemispherical domes to omega-shaped pits during CME at high tension^12^. However, it has been puzzling how cells overcome the transition from flat membrane to hemispherical domes during CME at high tension. Here we uncovered epsin’s ENTH and IDP domains play complementary roles to ensure the completion of CCS maturation under high tension environments (Fig. 7). Under a SIM-TIRF field, a productive CCS track shows evolution of the hemispherical dome manifested as a ring^30^. We found that overexpressing epsin or epsin EGFP in cells support the maturation of coated pits at high tension. Utilizing dual-color TIRF imaging, we showed that epsin EGFP recruitment into CCSs increases with an increase in resting membrane tension or acute membrane tension. We showed that masking ENTH domain activity in epsin (i.e. by placing EGFP at the N-terminus end of epsin) blocks the early recruitment of epsin to the CCS sites. The IDP domain provides binding to AP2 and clathrin, which is required for epsin’s ability to stabilize CCSs, since ENTH domain alone is cytosolic. Our finding is significant as most membrane-associated proteins disassemble or dissociate from the lipid bilayer as membrane tension increases^22,35,41^. The complementary actions of ENTH and IDP domains of epsin enable it to detect membrane tension variations to support the flat-to-dome transition in high tension environments. Previously, Brady *et al*. showed that both ENTH and C-terminal domain of epsin regulate its dynamic interaction with CCSs in *Dictyostelium*^42^. In contrast, our experiments showed that the C-terminus IDP domain that contains AP2 and clathrin binding sites are sufficient for epsin to target to CCSs. Further, Zeno *et al*. found that disordered domains enhance curvature sensitivity of structured domains^37^. Similarly, we find the synergy between a structured (i.e. ENTH) domain and an unstructured (i.e. IDP) domain can achieve membrane tension sensing^37^.

**Figure 7.**
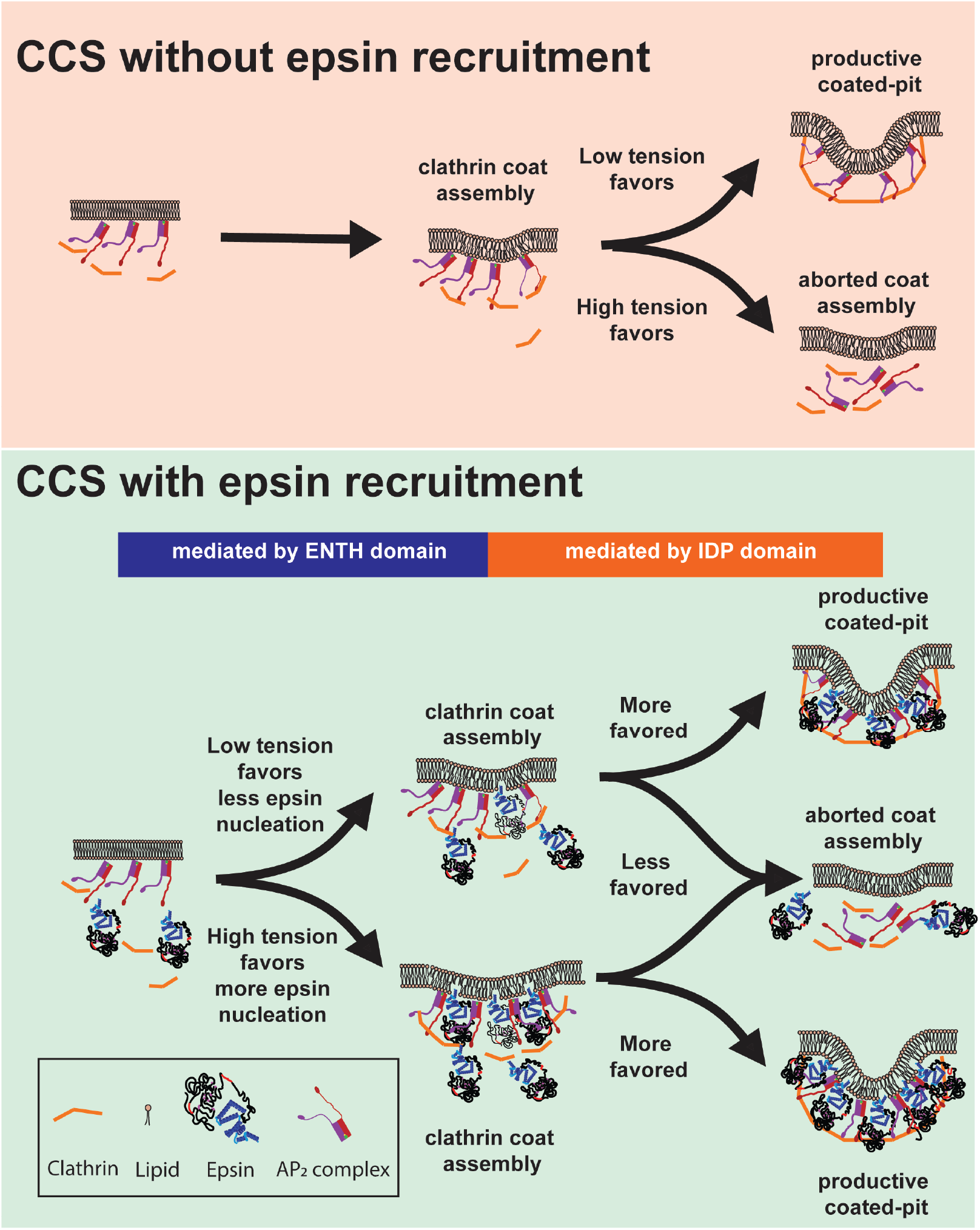
Summary of the role of epsin in CCS formation under different tension environments. Epsin provides enhanced CCS stability compared to CCSs without epsin. At high tension, more epsin molecules are recruited to counteract elevated tension and to provide stability to CCSs. The AP2 and clathrin binding of epsin primarily impart the additional stability to coated pits by anchoring constituents of CCS complex together.

The increase of epsin recruitment at high tension is dependent on ENTH domain, and specifically the H_0_ helix. Our observation that undisrupted activity of ENTH, specifically the amphipathic insertion of H_0_ helix is required for early epsin recruitment would be consistent with its tension-sensing role. Importantly, it is consistent with a previous finding where epsin recruits clathrin to PIP_2_-containing lipid monolayer and stimulates clathrin lattice assembly^15^. When ENTH of epsin is masked by N-terminal EGFP, its recruitment to CCS sites synchronizes with clathrin, presumably by binding to AP2 *via* epsin’s IDP domain. This is in agreement with our finding that removing the ENTH domain did not inhibit the recruitment of epsin to CCSs but only affected its early recruitment. Because IDP domain contains multiple clathrin and AP2 binding sites, it is likely the stability of CCSs at high tension is afforded by the avidity provided by multiple interactions^7^. The importance of multiple interactions is reinforced by the finding that knocking down either CHC or α-adaptin renders full-length epsin cytosolic. Although epsin ΔENTH can be recruited to sites of CCSs, this recruitment is reduced drastically when tension is increased, similar to the behaviors of other membrane-associated proteins when encountering high tension.

Using MD simulation, we showed that insertion of the H_0_ helix into the lipid bilayer enables recruitment of ENTH into the membrane. The simulations also show that membrane-association triggers ordering of H_0_ into a helical structure. Because of the favorable Ho–lipid interactions, ENTH prefers to bind to the membrane. Also, the H_0_ helix insertion displaces lipids, thereby providing an energetic advantage for epsin recruitment in high membrane tension regime. Tension-induced amphipathic helix insertion could perhaps be a general mechanism for membrane tension sensitivity for other cellular machineries. Under load, actin networks become denser to support the increased load^43^. In this context, increased membrane tension should promote actin assembly, and this can be achieved by increasing actin filament nucleation. Interestingly, nucleation promoting factor N-WASP has an amphipathic helix^44^. Thus, it is plausible, although highly speculative, that recruitment of N-WASP to the plasma membrane could be tension-sensitive in a fashion similar to epsin.

Although the H_0_ helix can insert into the membrane and acts as the tension sensor, our data does not support ENTH alone as the curvature generation machinery. We believe the ability of the IDP domain to tether different endocytic proteins (AP2, clathrin), besides generating steric pressure, is partly responsible for the stabilization effect of CCSs under high tension (i.e., epsin-positive CCSs have a larger fraction of productive structures). We speculate that steric repulsion by the bulky IDP domain also plays a role in inducing and stabilizing the curvature of the dome structure^16,17^. While stable epsin recruitment to CCSs requires the presence of CHC and AP2, once epsin is recruited to CCSs it supports the formation of CCS dome with the help of its IDP domain. Interestingly, cells overexpressing epsin-EGFP have more productive/stalled CCSs under high tension compared to cells overexpressing epsin. The EGFP following the IDP domain increases its bulkiness and therefore steric pressure. Our work implies the existence of a bi-directional stabilization between endocytic components of CCSs and IDP-mediated curvature generation.

Although we have identified a new mode of epsin function, we believe epsin is not unique in mediating tension-responsive recruitment to and stabilization of CCSs. In particular, adaptor protein 180 (AP180) and its homolog clathrin assembly lymphoid myeloid leukemia protein (CALM) both have alpha helices in their N-terminal domains^45^ along with an IDP domain consisting of endocytic binding domains^46^. Thus, it is plausible that a similar mechanism could be at play with AP180 and CALM to support CME under high tension environments. Several works using MD simulation and *in vitro* experiments have shown the role of amphipathic helices, specifically H_0_ helix, in curvature generation mechanism of N-BAR proteins^20,36,47,48^. It should be investigated whether H_0_ helix insertion of these proteins also enables tension-mediated membrane curvature generation and stabilization. Our work also provided some unexpected findings for future investigation. While we mainly focused on the role of epsin in high tension environments, decreasing tension in hypertonic solutions resulted in distinct responses. In particular, the initiation density and steady state density of epsin ΔENTH puncta under a hyper-osmotic condition was significantly higher compared to the iso-osmotic condition and to the full-length epsin under a hyper-osmotic condition. Furthermore, recruitment of epsin ΔENTH or epsin mut-H0 was significantly elevated under hyper-osmotic conditions. Interestingly, there appeared to be more clusters of CCSs in our SIM-TIRF images under hyper-osmotic conditions. It has been suggested recently that lowering membrane tension increases caveolar cluster formation^49^. Although clusters of CCSs have been postulated to be non-terminal endocytic events^50^, it will be intriguing to see if these represent local hot spots where membrane tension is low.

Finally, how might epsin’s recruitment to CCSs under high tension be important to cell physiology? Changes in cell tension is expected to trigger various cell signaling responses. Epsin has a ubiquitin-interacting motif (UIM) with the dual function of binding ubiquitin and promoting ubiquitylation. Thus, regulating epsin recruitment to CCSs has important implication of ubiquitinated cargo endocytosis^51^ and signaling. The association of epsin with ubiquitinated cargo is negatively regulated by clathrin^52^. An increase in resting cell tension increases epsin recruitment and decreases clathrin recruitment. Coupled with the increased stability of epsin-positive CCSs, our findings would suggest high membrane tension would impact ubiquitinated cargo endocytosis and signaling. The intersection of mechanosensing and endocytic regulation remains an under-studied area that will have profound implications in mechanotransduction.

## Methods

### Cell culture

Retinal pigment epithelial (RPE) cell was a gift from Sandra Schmid (UT Southwestern Medical Centre). The cells were maintained in Dulbecco’s Modified Eagle Medium with nutrient mixture F-12 (DMEM/F12) supplemented with 10% (v/v) fetal bovine serum (FBS) and 2.5% (v/v) penicillin/streptomycin at 37°C and 5% CO_2_.

### Generation of epsin EGFP and EGFP epsin lentiviral constructs

All constructs generated for this work have been sequenced for accuracy. EGFP epsin from pMIEG3 vector (gift from Sandra Schmid, original construct from JoAnn Trejo, UCSD) was amplified with the following primers (Integrated DNA Technologies Inc.) and cloned into pLVX puro vector, using BsrGI and Xbal digestion: Forward primer - 5’- ATAATATGTACAAGTCCGGACTCAGATCTC - 3’; Reverse primer - 5’ – GCGGCGTCTAGATTATAGGAGGAAGGGGTT - 3’. Epsin is amplified from pLVX puro vector, using the following primers and cloned into pLVX EGFP puro vector, using XhoI and EcoRI digestion to create epsin EGFP construct: Forward primer 5’ – CATCATCTCGAGGCCACCATGTCGACATCATCGCTG - 3’; Reverse primer - 5’- ATAATAGAATTCTCCACTTCCACTTCCACTTCCTAGGAGGAAGGGGTT -3’. Both constructs were transformed into competent *E. coli* cells (New England Biolabs Inc.) and selected using antibiotics (Ampicillin).

### Generation of epsin mutant constructs

Epsin mutant constructs were generated successively by mutagenesis using Q5^®^ Site-Directed Mutagenesis Kit (New England Biolabs Inc.). Clathrin binding domain 1 (CBD1) was deleted using following primers to create epsin ΔCBD1 EGFP: Forward primer 5’- TTCACAACCCCAGCCCCT-3’; Reverse primer 5’-AGATGACTCCTCCTTGCCC-3’. Clathrin binding domain 2 (CBD2) is deleted from epsin EGFP and epsin ΔCBD1 EGFP using following primers to create epsin ΔCBD2 EGFP and epsin ΔCBD 1 & 2 EGFP: Forward primer 5’- TCACTGGTGAGCCGACCA-3’; Reverse primer 5’-GGCTGCATTAGGGCCTAG-3’. Epsin amino acid sequence containing endocytic binding cites CBD1, CBD2 and AP2 binding motifs were deleted using following primers using to create epsin Δ EBD EGFP: Forward primer 5’- TCACTGGTGAGCCGACCA-3’; Reverse primer 5’-AGATGACTCCTCCTTGCCC-3’. ENTH domain of epsin was removed using following primers to generate epsin Δ ENTH EGFP: Forward primer 5’-GCCCACGCGCTCAAGACC-3’; Reverse primer 5’- CTTCATCTGACGCCGCAGC-3’. IDP domain of epsin was removed using following primers to generate epsin ΔIDP EGFP: Forward primer 5’- GGAAGTGGAAGTGGAAGTGGAGAATTCG-3’; Reverse primer 5’- GCGCTCCTCCCGAAGCCG-3’. H_0_ helix in ENTH domain of epsin was substituted to alanine using following primers to generate epsin mut-H0 EGFP: Forward primer 5’- CACCACCACCACACAAGCAATGATCCCTGG-3’; Reverse primer 5’- ATGATGATGATGCTCTGAGTAGTTGTGGAC -3’. All mutant constructs were transformed into competent *E. coli* cells (New England Biolabs Inc.) and selected using antibiotics (Ampicillin).

### Lentivirus transduction

Lentiviral vectors encoding epsin or epsin mutants were generated by transfecting 70% confluent HEK 293T for 24 h with a plasmid cocktail. The plasmid DNA cocktail contained 1.875 μg psPAX2, 625 ng pMD2.G, 2.5 μg pLVX vector diluted in 3 mL Opti-MEM (Thermo Fisher) containing 15 μL Lipofectamine 2000 (Thermo Fisher) and 5 mL of DMEM media with 10% FBS (v/v). The plasmid cocktail was replaced with fresh DMEM media containing 10% FBS after 24 h. The supernatant was harvested after 48 h, filtered through 0.3 μm sterile filter and flash frozen using liquid nitrogen.

### Stable cell line generation

RPE cells stably expressing EGFP-tagged epsin or epsin mutant and red fluorescent protein (mCherry)-tagged clathrin light chain a (mCherry-CLC) were generated in a two-step process. RPE cells were transduced with retroviruses (encoding mCherry-CLC) in a pMIEG3 vector produced by the UM vector core, followed by FACS sorting (UM Flow Cytometry core) to generate stable RPE cells expressing mCherry CLC. RPE cells expressing mCherry CLC were further infected with lentivirus encoding EGFP-epsin or epsin mutants followed by antibiotic selection (Puromycin) to generate double stable RPE cells.

### shRNA knockdown

shRNA constructs for epsin/alpha adaptin/clathrin heavy chain (CHC) was created in pLKO.1 vector by the UM vector core. Lentiviruses encoding shRNAs of interest were generated using the aforementioned lentivirus transduction method. RPE cells were infected with the viruses and selected using antibiotic selection (Puromycin). All downstream assays (TIRF imaging or Western blotting) were performed on day 5 of transduction and the level of knockdown was confirmed by Western blot analysis.

### Cell spreading on fibronectin islands

Polydimethoxysiloxane (PDMS) stamps with square shapes of size 25 μm and 32 μm were created from a silicon master mold made by using soft lithography. Sylgard-184 elastomer and curing agents (Dow Corning, Midland, MI) were mixed at a ratio of 10:1 (w/w) and casted over the silicon mold and cured at 60 °C overnight. Fibronectin (Sigma) solution (40 μg/ml) was added onto the stamps and incubated for 1 hour at room temperature. The stamps were blown dry using filtered air and placed in conformal contact with UV-ozone-treated PDMS-coated coverslip. Coverslips were spin-coated with a layer of PDMS diluted in hexane (1:20) at 5,000 rpm for 2 minutes. PDMS-coated coverslips enable efficient transfer of stamped proteins. Immediately after stamping, the coverslip was passivated with 0.1% (v/v) Pluronic-F127 (Sigma) for 1 hour, followed by extensive washing with PBS. RPE cells were seeded on the coverslips and allowed to selectively adhere to the square patterns. After 1 hour of seeding, media was changed to remove non-adhering cells. The adherent cells were allowed to fully spread for 5 hours, followed by TIRF imaging.

### Osmotic shock

The RPE cells were imaged in three media having different osmolarities: (i) hyper - 440 mmol/kg, (ii) iso – 290 mmol/kg, (iii) hypo – 220 mmol/kg) were used. Hyper-osmotic solution was prepared by adding 150 mM sucrose to phenol red-free DMEM containing 2.5% FBS. Iso-osmotic solution was DMEM media with 2.5 % FBS. Hypo-osmotic solution was prepared by adding deionized water containing 2.5% FBS to DMEM media containing 2.5% FBS in 1:3 ratio. All media osmolarities were checked on a Vapro osmometer. The cells were imaged between 5 minutes and 35 minutes of adding the different media.

### Confocal microscopy

The 3D profiles of cells subjected to different osmotic conditions were imaged using Olympus-IX81 microscope with spinning disk confocal scanner unit (CSU-X1; Yokogawa, Japan), EMCCD camera (iXon X3; Andor, South Windsor, CT), 60× objective (NA = 1.42). A z-step size of 0.2 μm was used. EGFP was used as volume marker. The 3D reconstruction was performed using 3D projection plugin in ImageJ.

### Micropipette aspiration

Variation in membrane tension due to osmotic shock was quantified by using micropipette aspiration. Glass micropipettes of inner diameter of ~5 μm were fabricated by pulling borosilicate glass pipette (BF100-50-10; Sutter Instrument) using a micropipette puller (Sutter Instrument). The micropipette was attached to a custom-made stage with pipette holder assembly (MI-10010; Sutter Instrument). An open chamber was made on coverslip seeded with RPE cells using VALAP sealant. The cells were subjected to osmotic shock, using aforementioned method. The micropipette was made into contact with the cells with the aid of brightfield imaging using an inverted microscope (Nikon TiS) equipped with 20 x objective and a CCD camera (CoolSNAP MYO). A negative hydrostatic pressure of 2.156 kPa was applied to the cell to aspirate the plasma membrane of cells into the pipette. The ratio of equilibrium protrusion length of membrane (*Lp*) vs inner radius of pipette was measured for different osmotic conditions.

### Live cell imaging *via* total internal reflection fluorescence (TIRF) microscopy

RPE cells expressing the constructs of interest were plated on a coverslip at a low concentration (~ 1.7 x 10^5^ cells per 22 x 22 coverslip) and allowed to spread for 12 to 16 hours. TIRF microscopy was performed using a Nikon TiE-Perfect Focus System (PFS) microscope equipped with an Apochromat 100X objective (NA 1.49), a sCMOS camera (Flash 4.0; Hamamatsu Photonics, Japan), and a laser launch controlled by an acousto-optic tunable filter (AOTF). Cells were imaged at 2 s intervals (100 ms exposure) for 5 min at 37 °C with dual-color excitation of 488 nm and 561 nm lasers (Coherent Sapphire).

### Image analysis for CCS dynamics

Image analysis was performed using CMEAnalysis software developed by Aguet *et al*^27^. mCherry channel was assigned as the primary detection channel and EGFP as the associated secondary channel. The program uses Gaussian mixture model fitting to detect and localize CCSs. It also performs CCS tracking using μ-track package with a gap-closing feature generating trajectories of CCSs. Only CCSs with a lifetime between 10 s and 120 s were considered for the downstream analysis. The program generates the lifetime and intensity data for CCSs under two categories (i) CCSs containing EGFP-epsin and (ii) CCSs not containing EGFP-epsin. These were further classified into six cohorts according to their lifetimes (cohorts (10–18, 20–38, 40–58, 60–78, 80–98, and 100–120 s). For quanitification of early arrival of epsin, aforementioned analysis was repeated with EGFP as primary detection channel and mCherry as the associated secondary channel.

### Live cell imaging *via* structural illumination microscopy in total internal reflection fluorescence mode (SIM-TIRF)

RPE cells expressing the constructs of interest were plated on a MatTek dish at a concentration of ~ 1.7 x 10^5^ cells per dish for 12 to 16 hours. SIM-TIRF was performed using a Nikon N-SIM microscope equipped with an Apochromat 100X objective (NA 1.49) and a sCMOS camera (Flash 4.0; Hamamatsu Photonics, Japan). Epsin EGFP was imaged using 488 nm laser with exposure time of 200 ms, with nine images taken in TIRF mode with linear translation of Moire pattern for SIM reconstruction. Similarly, mCherry clathrin was imaged using 561 nm laser with exposure time of 500 ms. Time interval between each set of SIM reconstruction images were 5 sec. Dual color SIM-TIRF images for epsin EGFP and mCherry clathrin was performed using DeltaVision OMX SR system (GE) equipped with 60x 1.42 NA objective and a sCMOS camera. Epsin EGFP and mCherry clathrin were imaged using 488 nm and 561 nm lasers respectively, with exposure times of 50 ms. Nine images were taken in ring TIRF mode with angular translation of Moire pattern for SIM reconstruction.

### Image reconstruction and analysis for SIM-TIRF

Images captured in Nikon N-SIM system was reconstructed into SIM images using Nikon Elements software. Reconstructed images were further equalized in intensity across time period using Nikon Elements software. Clathrin-coated structures (CCSs) were detected and tracked using Trackmate plugin in FIJI (ImageJ)^53^. A detector with Laplacian of Gaussian Filter is applied to detect CCSs with a quadratic fitting scheme for subpixel localization and estimated blob size parameter of 500 nm. A simple Linear Assignment Problem (LAP) tracker was applied with a maximum linking and gap closing distance of 500 nm and maximum gap closing of 2 frames. CCSs were morphologically characterized into abortive, productive and stalled structures.. Using lifetimes limits for abortive, productive and stalled CCS, the fraction of CCSs belonging to each category was determined.

### Molecular dynamic simulations

The MD simulation systems were composed of a bilayer with an ENTH domain of epsin (PDB number 1H0A^15^), a PIP_2_ molecule and 99 POPC lipids in the top leaflet and 100 POPC molecules in the bottom leaflet. In addition, the solution consisted of TIP3 (an all atom model of water) water molecules with 0.15 mM KCl. The simulations were performed in GROMACS using 303.15 K and 1 bar using CHARMM36 force field^54^.

The file taken from the protein data bank contained an ENTH domain of epsin and a PIP_2_ head in the inserted configuration. The head group was replaced with the entire lipid preserving the position and original orientation of the head group. Then, the POPC molecules were inserted in a grid pattern in order to construct the bilayer. Finally, the solution with the corresponding KCl concentration was added. All the files detailing the molecule and the equilibration procedure were obtained from CHARMM GUI website^55^.

In order to compute different positions of ENTH with respect to the membrane, we pulled the protein sequentially starting from the inserted configuration by imposing a restraining force on the center of mass of the protein. Additional restrains were applied to the PIP_2_ lipid to prevent it from leaving the membrane. The pulling proceeds until the protein is not in contact with the membrane. All protein configurations were given an initial equilibration time of 70 ns and a production run of 150 ns.

The secondary structure analysis tool of VMD^56^ was applied on the last 100 ns of production run to obtain the secondary structure for each residue in a given frame. The helicity plot was subsequently created by calculating the ratio of frames classified as a helix divided by the total number of frames for each residue.

The area plot for the inserted proteins were obtained using g-lomepro^57^. We used 100 ns production run after an initial equilibration of 100 ns. Since the protein has no restrain of movement along the plane of the bilayer during the simulation, the frames had to be centered around the H_0_ helix center of mass.

### Transferrin uptake assay

RPE cells expressing epsin mutants were serum-starved for 4 h. Cells were subjected to hypo- and iso-osmotic shock for 10 min and then allowed to uptake transferrin Alexa 647 (25 μg/ml) (Thermo Fisher) for a further 10 min, followed by immediate fixation with 4% paraformaldehyde (Electron Microscopy Sciences) in PBS for 10 min.

### Western blot

RPE cells were lysed with RIPA buffer (Thermo Fisher) containing protease inhibitor on ice for 10 min. 25 μL of lysate mixed with sample buffer (1:1 ratio) (Bio-Rad) was loaded per lane in a 10% SDS-PAGE gel (Bio-Rad). The proteins were transferred to nitrocellulose membrane and blocked for 1 h with 3% BSA solution (in PBS). Primary antibodies against epsin (Abcam: ab75879 (1:500 dilution)), CHC (Abcam; ab2731 (1:500 dilution)), α-adaptin (Abcam: ab2807 (1:100 dilution)) were used to quantify the expression levels of the respective proteins in epsin overexpressed cells, epsin/CHC/α-adaptin knockdown cells and wild type cells. Blots with primary antibody and 1:1000 dilution of anti-GAPDH antibody were incubated overnight followed by 1 h incubation of secondary antibody conjugated to Dylight 680 nm or Dylight 800 nm. The blots were imaged using an LiCor imaging system or an Azure imaging system.

### Statistical analysis

For intensity, lifetime and initiation density data, the statistical significance was verified by oneway ANOVA test. Subsequently, two-tailed student t-test was performed between pair of data to determine the *p*-values. *, **, and *** was assigned to *p* < 0.01, *p* < 0.001 and *p* < 0.0001 respectively.

## Data availability

The data that support the findings of this study are available from the corresponding author upon request. The computational code can be made available upon request to A.A.

## Acknowledgements

This work was supported by the National Science Foundation NSF-MCB 1561794 to A.P.L. and NSF grants CMMI 1562043 and CMMI 1727271 to A.A.. We acknowledge Dominic Ciarelli for helping us with micropipette aspiration, Aaron Taylor and Eric Rentchler from UM microscopy core for helping us with TIRF-SIM imaging.

## Author contributions

J.G.J., A.A. and A.P.L. conceived the study, J.G.J., C.O., A.A., A.P.L. designed the experiments, J.G.J. and V.Y. performed the experiments, C.O. carried out the molecular dynamics simulation, J.G.J, A.A., and A.P.L. wrote the manuscript. All authors commented on the manuscript and contributed to it.

## Competing interests

The authors declare no competing interests.

