## Supplemental figures for "Complimentary action of structured and unstructured domains of epsin supports clathrin-mediated endocytosis at high tension"

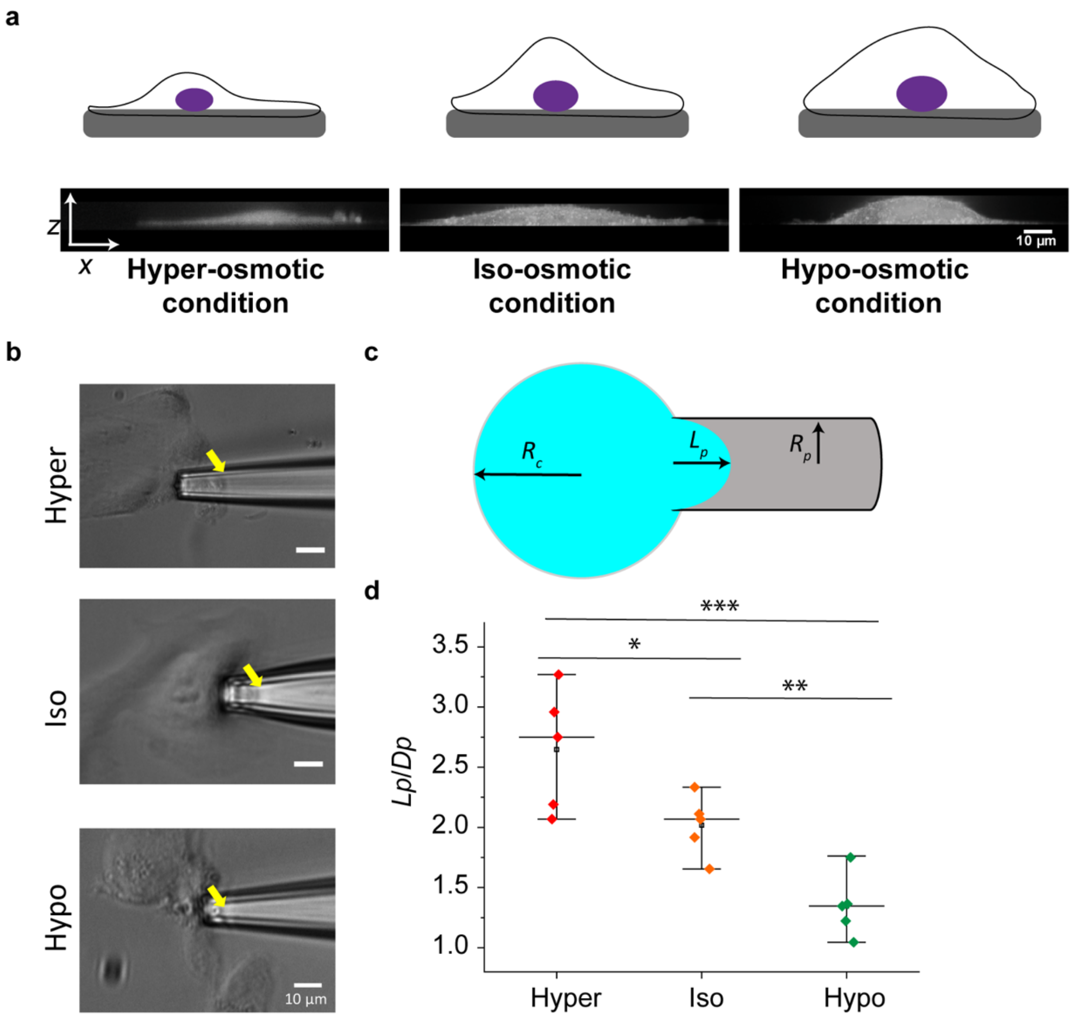


**Supplementary Figure 1. Micropipette aspiration shows osmotic shock increases tension. a.** Osmotic shock experiment for acute tension drop and spike, as described in the Materials and Methods. Representative confocal cross section images reconstructed from 3D image stacks of cells under hyper-, iso- and hypo-osmotic conditions. **b.** Plasma membrane aspirated into the micropipette due to negative hydrostatic pressure ($\Delta P$ = -2.16 kPa) at hyper-, iso- and hypo-osmotic conditions. **c.** The simplified schematic of plasma membrane aspiration into the micropipette. $L_p$ is the total length of aspiration, $R_c$ is the radius of the cell (assuming spherical shape) and $R_p$ is the radius of the pipette ($D_p = 2R_p$). **c.** $L_p/D_p$ values of cells under hyper-, iso- and hypo-osmotic conditions. *, **, *** represent $p < 0.01$, $p < 0.001$, and $p < 0.0001$, respectively.

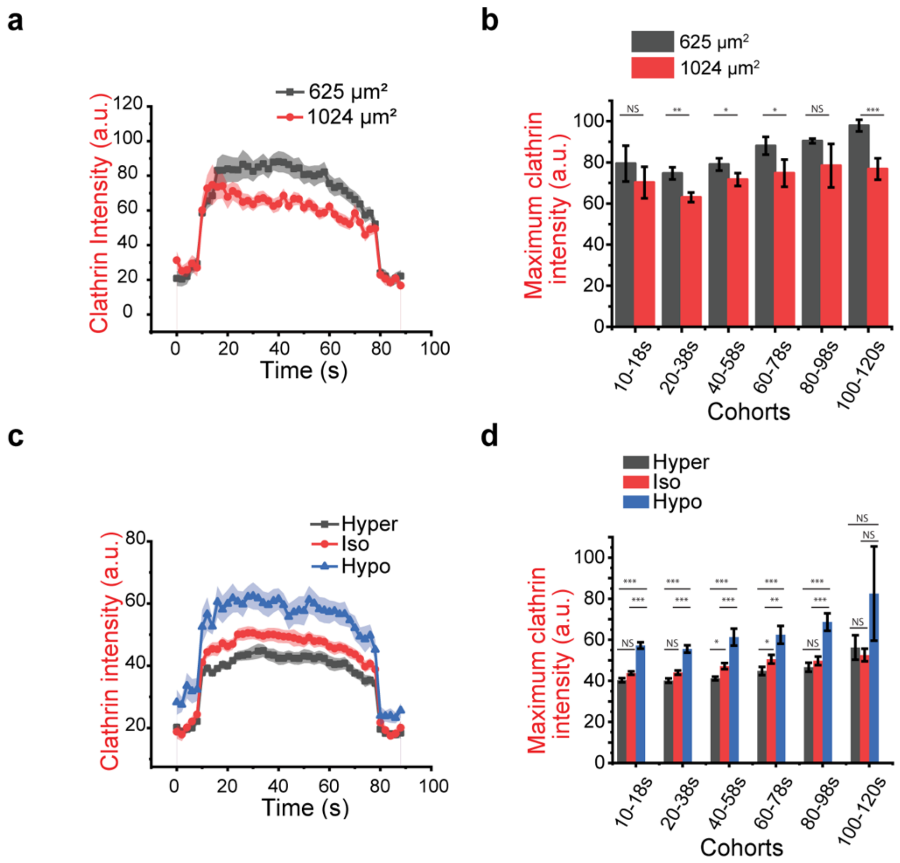


**Supplementary Figure 2. Clathrin recruitment into epsin-positive CCPs is distinct between increasing resting tension and acute tension increase. a.** The average intensity trace of clathrin in 25 (black) and 32 μm (red) islands for epsin-positive CCPs with 60 -78 s lifetime. **b.** The average plateau intensity of clathrin across different epsin-positive CCP lifetime cohorts. **c.** The intensity profiles of epsin-positive CCPs with 60 -78 s lifetime for different osmotic conditions. **d.** The average plateau intensity of clathrin across different epsin-positive CCP lifetime cohorts. For a and b, $n_{cells}$ for 25 μm square and $n_{cells}$ for 32 μm square are 18 and 19, respectively. For c and d, the number of cells for hyper-, iso-, and hypo-osmotic conditions were 19, 19, and 21, respectively. The error bars denote standard error. NS denotes not significant. *, **, *** represent $p < 0.01$, $p < 0.001$, and $p < 0.0001$, respectively.

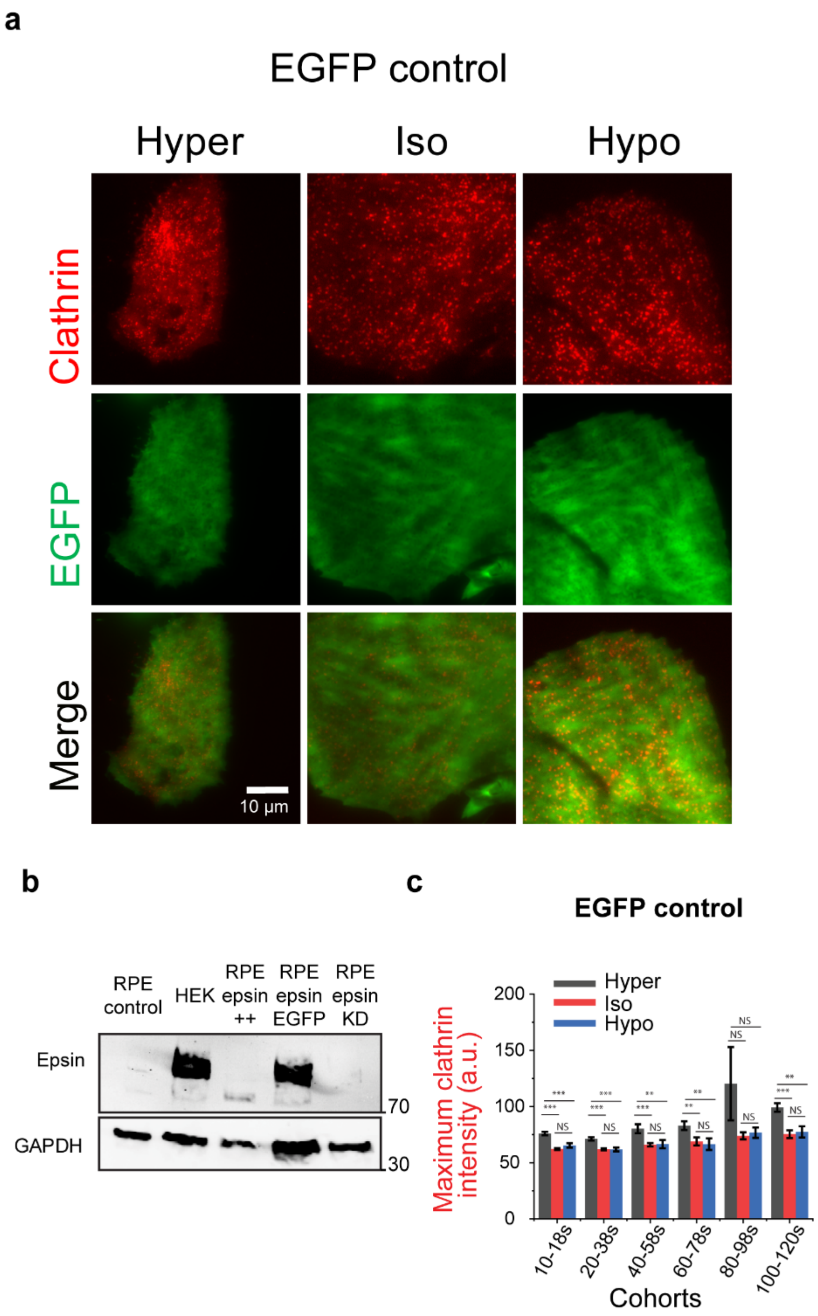


**Supplementary Figure 3. a.** Representative fluorescence images of clathrin and EGFP of RPE cells expressing EGFP as a control under different osmotic conditions. **b.** The average plateau intensity of clathrin across different CCP lifetime cohorts in RPE cells. For b the number of cells for hyper-, iso-, and hypo-osmotic conditions were 20, 19, and 16, respectively. The error bars denote standard error. **c.** Western blot showing the expression of epsin in RPE cells and HEK cells (highest epsin expression shown), RPE cells overexpressing WT epsin, epsin EGFP and knockdown of epsin, with GAPDH as a loading control. NS denotes not significant. **, *** represent $p < 0.001$, and $p < 0.0001$, respectively.

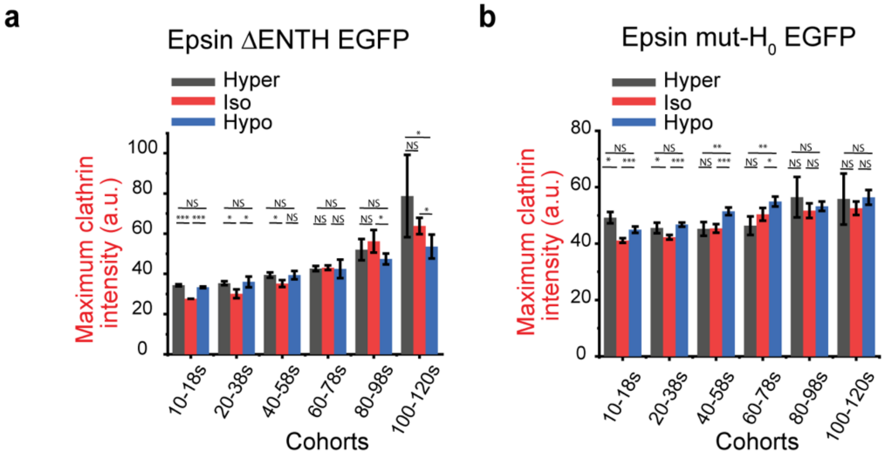


**Supplementary Figure 4. a.** The average plateau intensity of clathrin in different epsin ΔENTH-positive CCP lifetime cohorts in different osmotic conditions. **b.** The average plateau intensity of clathrin in different epsin mut-$H_0$ positive CCP lifetime cohorts in different osmotic conditions. For a, the number of cells for hyper-, iso-, and hypo-osmotic conditions were 20, 20, and 20 respectively. For b, the number of cells for hyper-, iso-, and hypo-osmotic conditions were 19, 24, and 18 respectively. The error bars denote standard error. NS denotes not significant. *, **, *** represent $p < 0.01$, $p < 0.001$, and $p < 0.0001$, respectively.

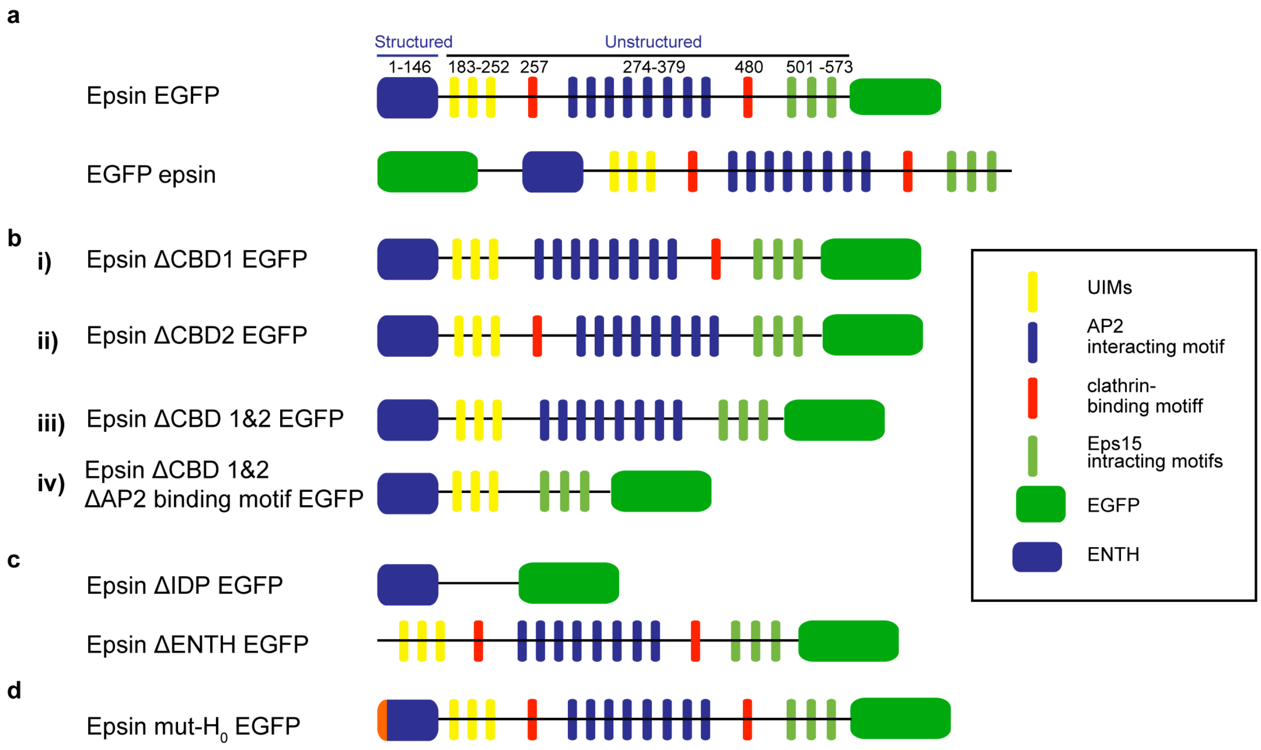


**Supplementary Figure 5. Epsin mutants. a.** Full length epsin with EGFP tagged on C-terminus and N-terminus. Different domains/motifs of epsin and their location is shown. **b**. Epsin EGFP mutants generated by deleting binding domains (i) clathrin-binding domain1 (CBD1), (ii) clathrin-binding domain 2 (CBD2), (iii) clathrin-binding domain 1 and 2 (CBD1 & 2), (iv) CBD1 & 2 and region containing repeated DPW motifs binding to AP2. **c.** Epsin mutant with entire unstructured region of the protein deleted (ΔIDP) and epsin with entire structured region of the protein deleted (ΔENTH). **d**. Epsin EGFP after mutating $H_0$ helix in the N-terminus of protein to alanine residues.

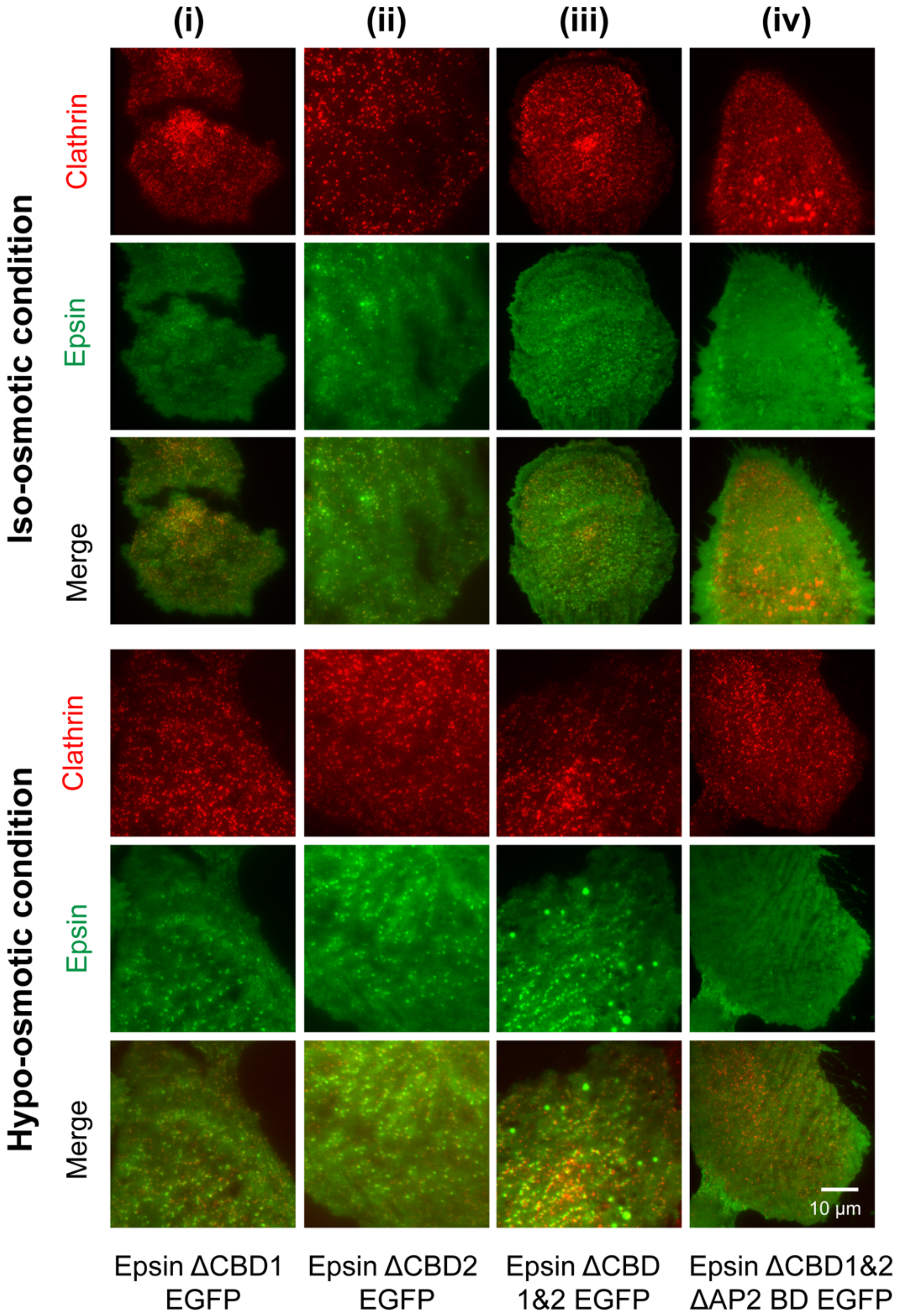


**Supplementary Figure 6. Removal of AP2 and clathrin binding sites render epsin cytosolic.** Representative fluorescence images of cells overexpressing EGFP-tagged epsin and epsin mutants under iso- (top panel) and hypo- (bottom panel) osmotic conditions**.** Cells overexpressing (i) epsin ΔCBD1 EGFP, (ii) epsin ΔCBD2 EGFP, (iii) epsin ΔCBD1&2 EGFP, (iv) epsin ΔCBD1&2 and ΔAP2 binding domains EGFP along mCherry CLC is shown.

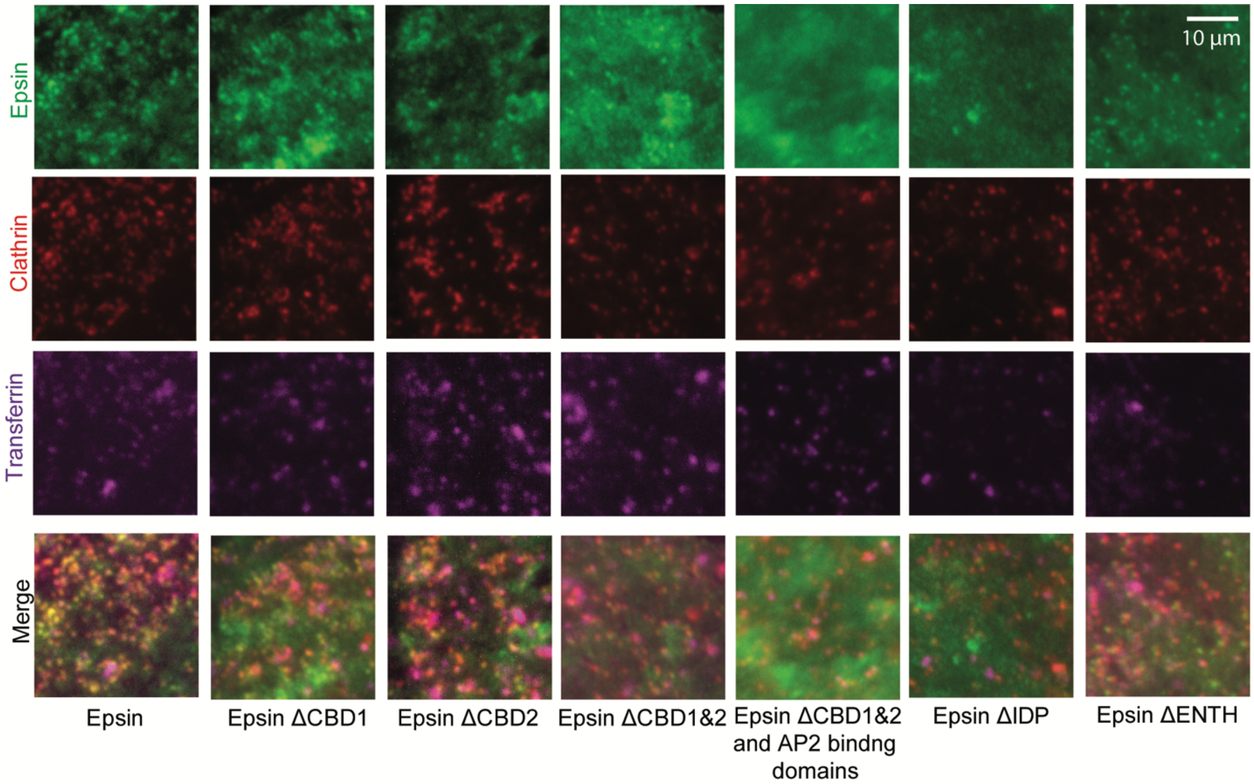


**Supplementary Figure 7. Transferrin uptake in cells overexpressing EGFP-tagged epsin and epsin mutants.** Representative fluorescence images of cells expressing epsin EGFP, epsin ΔCBD1 EGFP, epsin ΔCBD2 EGFP, epsin ΔCBD1&2 EGFP, epsin ΔCBD1&2 and ΔAP2 binding domains EGFP, epsin ΔIDP EGFP, epsin ΔENTH EGFP, along with mCherry CLC and Alexa Fluor 647 transferrin (25 µg/ml with 10 min incubation).

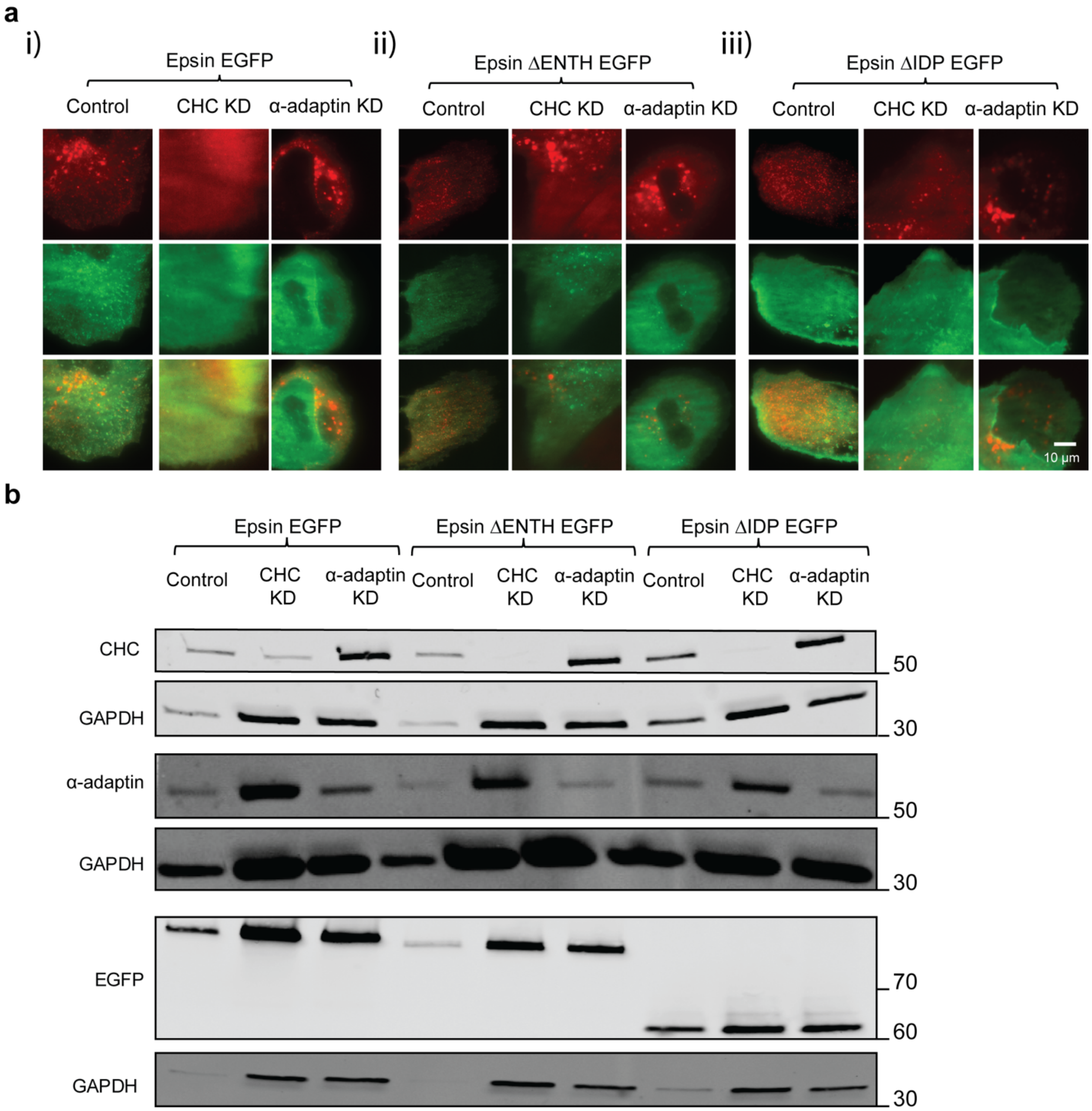


**Supplementary Figure 8. Knocking down clathrin heavy chain or AP2 subunit disrupts the recruitment of epsin**. **a.** Representative fluorescence images of (i) full length epsin EGFP, (ii) epsin ΔENTH EGFP and (iii) epsin ΔIDP EGFP, along with mCherry CLC in RPE cells with control (scramble shRNA), clathrin heavy chain (CHC) shRNA knockdown, and α-adaptin shRNA knockdown. **b.** Western blot showing the knockdown of CHC and α-adaptin, with GAPDH as a loading control. EGFP panel shows the expression of fusion protein epsin EGFP, epsin ΔENTH EGFP, and epsin ΔIDP EGFP.
